# Lepidopteran scale cells derive from sensory organ precursors through a canonical lineage

**DOI:** 10.1101/2024.05.31.596873

**Authors:** Ling S. Loh, Kyle A. DeMarr, Martina Tsimba, Christa Heryanto, Alejandro Berrio, Nipam H. Patel, Arnaud Martin, W. Owen McMillan, Gregory A. Wray, Joseph J. Hanly

## Abstract

The success of butterflies and moths is tightly linked to the origin of scales within the group. A long-standing hypothesis postulates that scales are homologous to the well-described mechanosensory bristles found in the fruit fly *Drosophila melanogaster*, where both derive from an epithelial precursor specified by lateral inhibition that then undergoes multiple rounds of division. Previous histological examination and candidate gene approaches identified parallels in genes involved in scale and bristle development. Here, we provide definitive developmental and transcriptomic evidence that the differentiation of lepidopteran scales derives from the canonical cell lineage, known as the Sensory Organ Precursor (SOP). Live imaging in moth and butterfly pupae shows that SOP cells undergo two rounds of asymmetric divisions that first abrogate the neurogenic lineage, and then lead to a differentiated scale precursor and its associated socket cell. Single-nucleus RNA sequencing across a time-series of early pupal development revealed differential gene expression patterns that mirror canonical lineage development, including Notch-Delta signalling components, cell adhesion molecules, cell cycling factors, and terminal cell differentiation markers, suggesting a shared origin of the SOP developmental program. Additionally, we recovered a novel gene, the POU-domain transcription factor *pdm3*, involved in the proper differentiation of butterfly wing scales. Altogether, these data open up avenues for understanding scale type specification and development, and illustrate how single-cell transcriptomics provide a powerful platform for understanding the evolution of cell types.

## Introduction

The colour patterns that adorn the wings of butterflies and moths consist of mosaics of hundreds of thousands of microscopic units known as scales. These cuticular extensions display an extraordinary diversity of colours and shapes, and ultimately form the building blocks behind the large pattern diversity of the insect order Lepidoptera, which is named after them (*lepis*, ancient Greek for “scale”). Their phenotypic diversity is largely linked to colouration and its functions in predator avoidance, sexual selection, and thermoregulation. Although they each emerge from a single cell and are limited to a thickness of 1-2 microns, scales have diversified to occupy the whole range of colour space, with pigments and nanostructures that can make them reflect or absorb specific parts of the ultraviolet and visible light spectrum (Thayer & Patel, 2023), produce some of the brightest or darkest known biomaterials (Davis et al., 2020; McDougal et al., 2019), or deflect radiative heat in the mid-infrared wavelengths (Tsai et al., 2020). In addition to their ability to interact with light, scales have also evolved a variety of other functions including pheromone emission, aerodynamic activity and acoustic camouflage (Gorb, 2001; Neil et al., 2020; Watson et al., 2017).

Scales are largely hollow, flattened chitinous structures, each secreted by a single scale-building cell during pupal development (Ghiradella, 2010; McDougal et al., 2021). Each scale-building cell is ensheathed by a socket-building cell, through which the base of the scale is anchored (Dinwiddie et al., 2014; Kristensen & Simonsen, 2003). Organised in ordered rows, scales cover the bilayered adult wing completely, with specific spatial arrangements to produce the final wing colour pattern (Köhler, 1932; Nijhout, 1991). Each scale acts like a pixel on a screen to build wing pattern, and the colour spectrum exhibited by each individual scale provides a readout of its underlying pigment composition and structural properties (Day et al., 2019; Gilbert, 2003; McMillan et al., 2020; Thayer & Patel, 2023). Overall, this makes scales a useful test case of phenotypic variation for studying how single-cell derived structures develop and diversify.

Scales relate to a more general class of chitinous extensions known as setae, which perform a wide range of functions in sensory reception, defense, colouration, and copulation across insects (Dickerson et al., 2021; Ghiradella, 1998; Keil, 1997; Richter et al., 2023; Tanaka et al., 2009; Tokunaga, 1962; Tuthill & Wilson, 2016; Winterton, 2009). Sensory bristles (sensilla) have been the most studied and share a stereotyped mode of development where each cell component derives from a series of asymmetric cell divisions (Hartenstein & Posakony, 1989; Lai & Orgogozo, 2004). In the fly *Drosophila melanogaster*, a typical sensillum is composed of a neuron, insulated by a sheath cell, and a shaft cell that forms the bristle accompanied by a socket cell surrounding the shaft (Hartenstein & Posakony, 1989). The four daughter cells, and occasionally a fifth glial cell that apoptoses shortly after its formation, are derived from a sensory organ precursor (SOP) via two successive rounds of asymmetric divisions (Fichelson & Gho, 2003; Lai & Orgogozo, 2004; Reddy & Rodrigues, 1999). The SOP first emerges within a proneural epithelium via processes of lateral inhibition and protrusion-mediated signalling, initiated by low-Notch/high-Delta expression prior to the prepatterning of bristles (Cohen et al., 2010; Corson et al., 2017; Simpson et al., 1999). Other factors that confer cells to proneural fates during SOP determination include proneural proteins containing the basic helix-loop-helix domain, like the Achaete-Scute complex (AS-C) (García-Bellido, 1979; Ghysen & Dambly-Chaudiere, 1988; Skeath & Carroll, 1994). Reviewing several decades of research, Lai and Orgogozo (2004) formalised the idea that all insect sensory organs are developmental variations on a common theme known as the SOP ‘canonical lineage’. *D. melanogaster* SOP derivatives follow a stereotyped pattern of division and differentiation, guided by a conserved gene regulatory network (Figure 1A). Variations from the canonical lineage result in morphologically distinct sensory organs, with modifications including lineage-specific cell proliferation or death, novel recruitment into the sensory cluster and the alteration of the terminal progenitor cell fates (Hopkins et al., 2023; Klann et al., 2021; Mangione et al., 2023).

**Figure 1:**
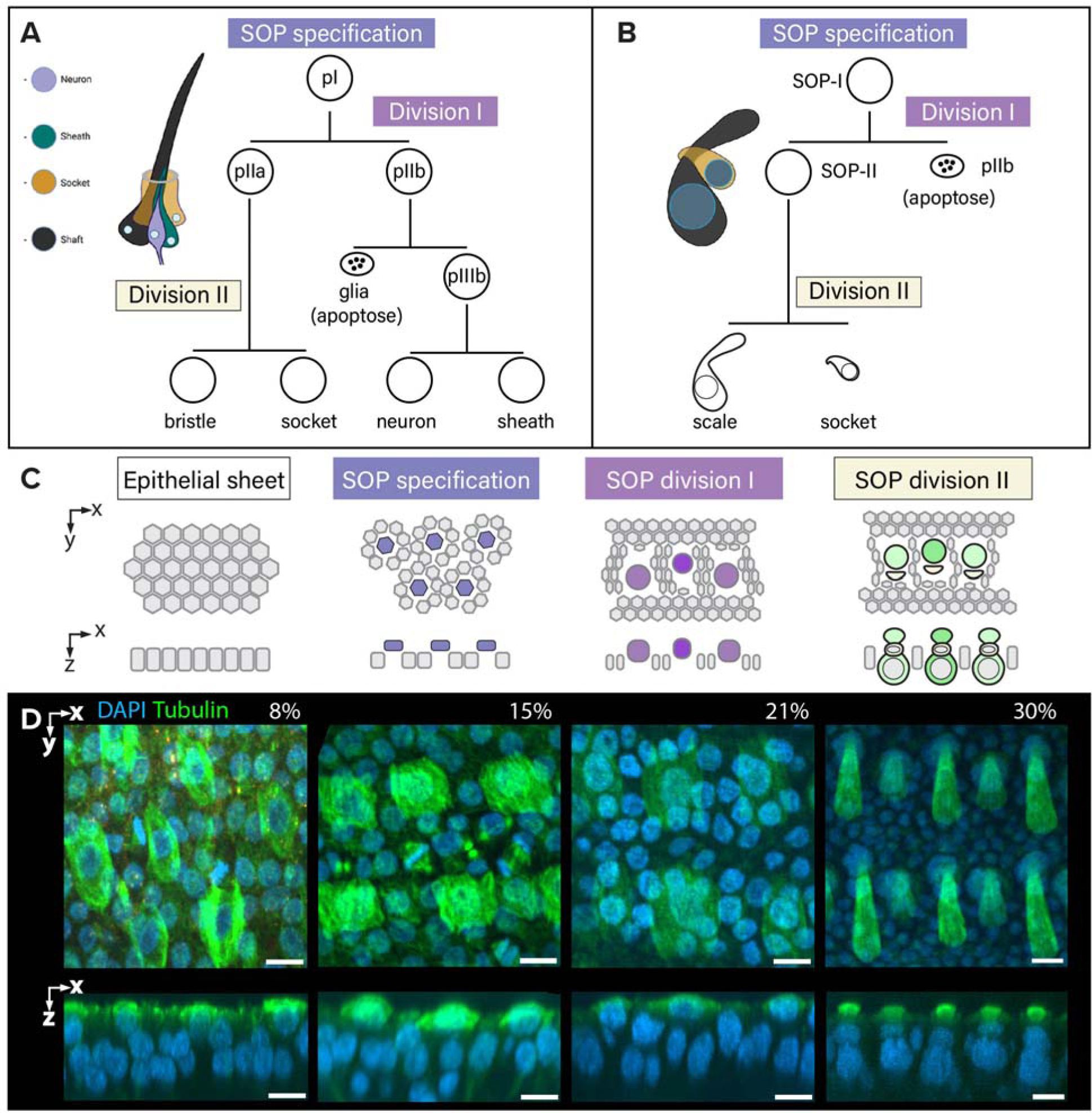
A visual summary of cellular events during SOP differentiation and division to produce fly mechanosensory bristles and lepidopteran scales. **A-B.** A comparative depiction of the canonical SOP lineage in *D. melanogaster* [A; adapted from Hopkins *et al*. (2023)], and the hypothetical lineage of lepidopteran wing scales (B), where pIIb and its derivatives are missing or not observed in histology and live imaging records (Ghiradella, 1998; McDougal et al., 2021; Stossberg, 1938). **C.** Model for the development of lepidopteran scale organs in a single-layered pupal wing epithelium. **D.** Anti-tubulin antibody (green) and DAPI (blue) stainings of *V. cardui* pupal wings showing representative stages of scale development. From 8-21% pupal development, tubulin is highly expressed in apically-localised SOPs and their derivatives, as well as in the mitotic epithelial cells. Following scale and socket cell differentiation (here seen at 30%, top layer nuclei = sockets; bottom layer nuclei = scale cell bodies), tubulin is remodelled into linear bundles within the secreted scale extensions. Scale bars = 10 μm. X-axis: antero-posterior; y-axis = proximo-distal; z = apico-basal.

Histological evidence of parallels between mechanosensory bristles and scales provided some support that scales are part of this ‘canonical lineage’. First, a similar ontogenetic series during SOP specification was described in the moth *Ephestia kuehniella* (Figure 1B; Köhler, 1932; Stossberg, 1938). Initially during pupal formation, the wing appears as a bilayered sheet of undifferentiated epithelial cells (Figure 1C). By around 8% pupal development, the cell homologous to the SOP has a large nucleus, which was reported in *Ephestia* to undergo a first asymmetric division perpendicular to the wing surface into an upper pIIa and a pIIb lower cell. The lower pIIb cell was observed to descend further and undergo apoptosis. The remaining hypothetical pIIa cell then undergoes a second asymmetric division into an upper socket-building cell and a lower scale-building cell (Stossberg, 1938). Each of the two rounds of asymmetric division occur in a wave along the proximodistal axis of the wing, and we refer to them as SOP-I prior to the first division and SOP-II prior to the second division into scale- and socket-building cells. Beginning around 15% pupal development, extracellular extensions start to emerge above the apical surface of the wing membrane, initially appearing as actin-filled tubular structures which then broaden into an oar-like shape (Dinwiddie et al., 2014). These scale extensions reach terminal lengths by 45% development, followed by the development of intricate surface ornamentation on the exposed surface by 60% pupal development (Figure 1D; Lloyd et al., 2024; Seah & Saranathan, 2023).

Similarities in histological changes between lepidopteran scales and mechanosensory bristles have prompted a few developmental studies of candidate genes involved in SOP lineage specification. The spatial definition of SOP clusters within the fly epithelium requires Notch (N) signalling to inhibit SOP fates (Troost et al., 2023). In early butterfly pupal wings, N is detected in rosettes of non-SOP epithelial cells, surrounding central SOP precursors that express low N (Reed, 2004). Loss of N increases SOP density, implying that N mediates SOP patterning via a lateral inhibition mechanism (Pomerantz, 2021). In flies, the activation of SOP fate in low-N cells requires the expression of transcription factor genes of the *achaete-scute* complex (Skeath & Carroll, 1991). Two lepidopteran gene homologues dubbed *ASH1* and *ASH2*, are transiently expressed in scale cell precursors, and *ASH2* is required for scale specification in both silkworms and butterflies (Galant et al., 1998; Pomerantz, 2021; Zhou et al., 2009), again suggesting homology of this process with flies. Finally, a recent single-cell transcriptome of *Bicyclus anynana* butterfly wings at 18% development showed that the transcription factor *shaven* (*sv*) is specifically expressed in scale cell precursors, and that its mosaic knockout results in scaleless clones (Prakash et al., 2024). This is identical to the described function of *sv* as a master specifier of the bristle shaft cell precursor in *Drosophila* (Fu et al., 1998). Altogether, the two lines of evidence from histological records and genetic work present the lepidopteran SOP specification and division into scale-building cells as strikingly similar to the canonical SOP lineage in the fly.

The remarkable precision of nanostructures and their replication across hundreds of thousands of scales on each wing surface speaks to the potential for lepidopteran scales as a system to understand the diversity and robustness of biological systems. However, we still have only a rudimentary understanding of how SOPs give rise to lepidopteran scale-building cells. To this end, we present here the first time-series dataset of single-nuclei resolution transcriptomes of a developing pupal wing. Nuclei were isolated from forewing tissues of the postman butterfly *Heliconius melpomene*, spanning a developmental time-series from 10% to 30% pupal development at 5% intervals, and an additional sample each for 10% and 25%. We also performed live imaging to identify the timing of SOP divisions in a cabbage looper moth *Trichoplusia ni* and the buckeye butterfly *J. coenia*, and tested marker genes in the pantry moth *Plodia interpunctella* and painted lady butterfly *Vanessa cardui*. This work serves to illuminate important aspects of cell fate specification and differentiation during wing scale development, decoding how self-patterning programs are modulated to ascribe a variety of cell types. Given the premise that a diverse repertoire of insect sensory organs stems from the modulation of a core process, we consolidate and address past observations to present the lepidopteran scale as a derivative of this canonical lineage.

## Results

### Live imaging reveals the early process of SOP specification

Serial dissections of tissue cannot resolve temporal dynamics of cell divisions, nor the developmental trajectory of a lineage; additionally, dissection of early wings from pupae can be technically difficult and compromises morphological features of the fragile tissue. Therefore, we employed a live imaging approach to document the series of cellular divisions leading up to the formation of the socket- and scale-forming cells in the cabbage looper moth, *Trichoplusia ni*, and the buckeye butterfly, *Junonia coenia* (Figure 2). The wings of newly eclosed pupae can be rearranged before the cuticle sclerotises, to allow visualisation through the transparent peripodial membrane of the hindwing and the introduction of Hoechst33342 allows the tracing of nuclear divisions through the fluorescent staining of DNA (Ohno & Otaki, 2015). Pupal wings were observed to be initially composed of uniform epithelial cells until around 5% development when the future SOP nuclei grew in size and neighbouring epithelial cells arranged into rosettes encircling these enlarged cells. SOPs in *T*. *ni* appeared to follow the canonical set of divisions described in *E*. *kuhniella*, with two obvious mitotic spindles forming and a degenerating pIIb cell appearing after the first division (Figures 2A, S1, <u>Movie S1</u>). By contrast, *J*. *coenia* SOPs appeared to undergo a singular complete division to form scale and socket daughters, seemingly foregoing a first division (Figures 2B, S2, <u>Movie S2</u>). Instead of a mitosis, the SOP nucleus appears to undergo partial DNA condensation, failing to form a metaphase plate, and then repositions itself without producing a degenerating pIIb daughter; subsequently, it settles into the epithelium and divides into the scale and socket. These results do not remove the possibility that an anucleate pIIb daughter cell is produced which would not be detectable with the Hoechst33342 stain.

**Figure 2.**
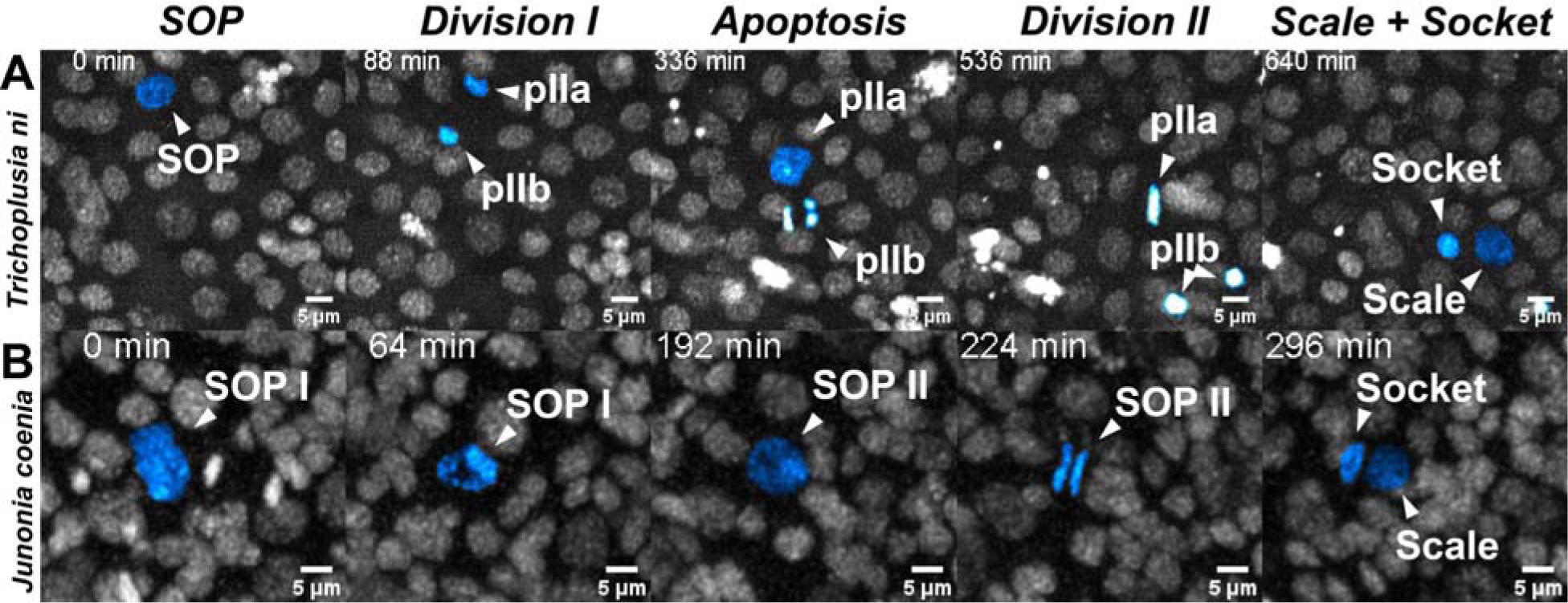
Live imaging of Hoechst33342 stained nuclei in *Trichoplusia ni* and *Junonia coenia* during early pupal wing development, from SOP-to scale/socket stages. **A.** Main cellular events highlighted in *T. ni* reveal the two rounds of mitotic division punctuated by an apoptotic event of the pIIb cell into two clusters of nuclear DNA aggregations (arrowheads in panel 4 ‘*Division II*’). Imaging of pupal hindwing began from 1h (0.7% development) to 24h APF (17% development). **B.** Main cellular events highlighted in *J. coenia* provide evidence for a mitotic division into scale- and socket-building cells. Imaging of *J. coenia* pupal hindwing began from 24h (13% development) through 40h APF (21% development). Scale bars = 5 μm. Abbreviations: APF = after pupa formation.

### A single cell time series of wings spanning early pupal developmental reveals cell type-specific marker genes

We employed an unbiased approach of single-nuclei RNA sequencing data (snRNAseq) on early pupal development. Seven pairs of forewings from five developmental time points (10%, 15%, 20% 25% and 30% pupal development) were obtained using *Heliconius melpomene*, beginning from the early phase of the growing SOP thorough to the period where the scale cell begins to rapidly extend (Figure 3A). We observed that large size of scales in the samples at 25% pupal development and onwards led to cellular debris inhibiting the recovery of GEMs. To compensate for this, we added a sucrose density gradient step during nuclei isolation for the two older samples (25%, 30% development). This may have indirectly enriched for denser polyploid nuclei of scale- and socket-building cells over the less dense non-polyploid epithelial and tracheal cells. After filtering for nuclei with less than 20% mitochondrial reads, we recovered the expression of 34,372 genes in 6,021 nuclei and identified six major clusters (Figure 3B-B’).

**Figure 3.**
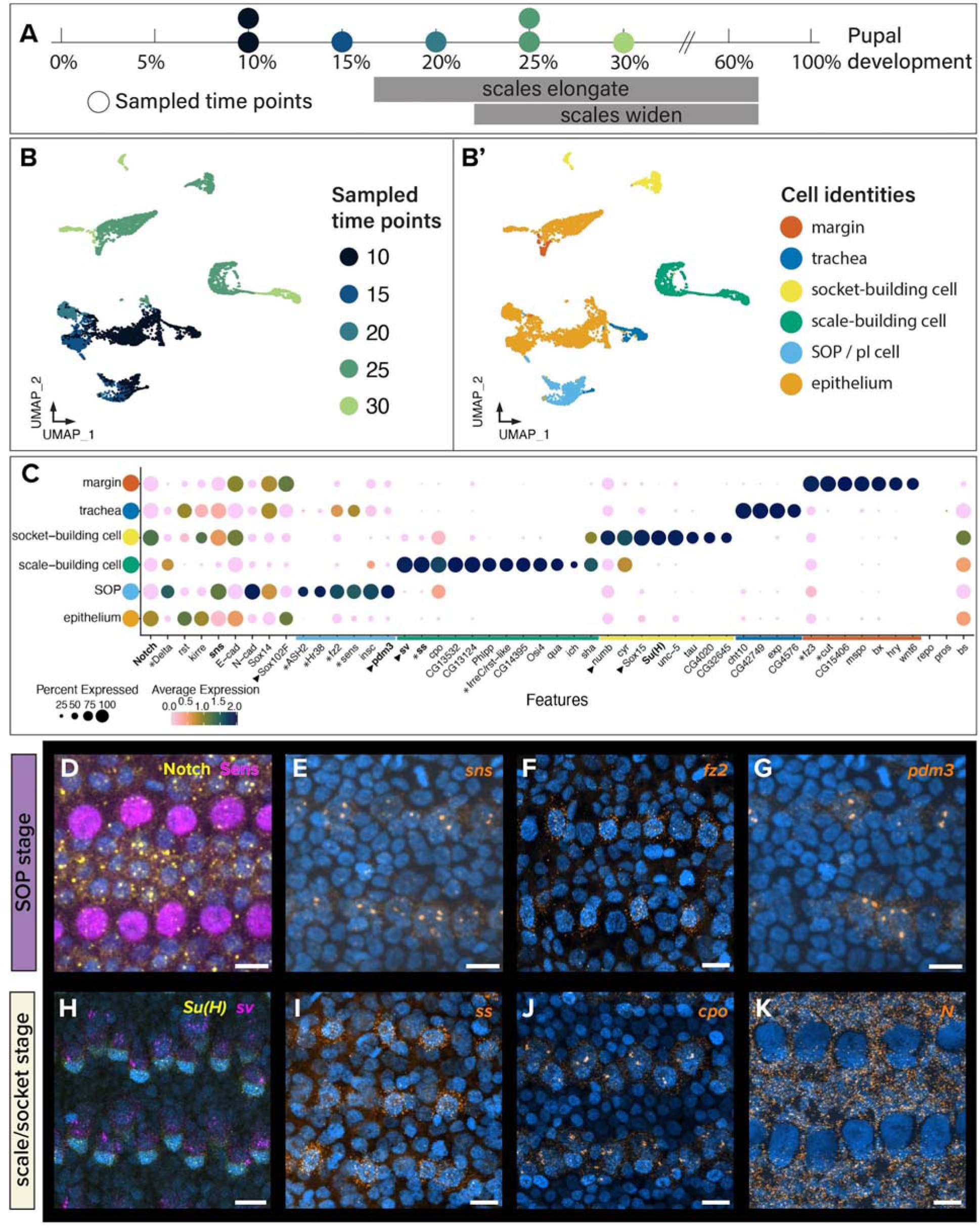
Cellular events and expression changes demarcate SOP- and scale/socket stages during 10-30% pupal development. snRNAseq data was obtained from *H. melpomene* and *in situ* expression of canonical marker genes were validated in *V. cardui*. **A.** Sampled timepoints coloured by legend in B. **B-B’.** Overall merged and integrated UMAP of snRNAseq data from seven pairs of *H. melpomene* forewings reveals six major cell types across five timepoints. **C.** Dotplot of putative marker genes with top expression in identified cell identities. Expression and/or functional data is present for genes with * (Banerjee et al., 2023; Hanly et al., 2023; Perry et al., 2016; Pomerantz, 2021; Prakash et al., 2024). Genes in bold were tested for expression in D-E’’’ and genes with black triangles were tested for function in Figures 5-6. **D-K.** Expression of selected marker genes using immunofluorescence (D: Sens and Notch protein) and hybridization chain reaction (HCR; E-K: mRNA detection of *fz2, pdm3, sns, Su(H)*, *sv, cpo*, *ss* and *N*) in *V. cardui* developing pupal wings during SOP (19-23 h APF, 11-14% development) and scale-/socket-building stages (27-48 h APF, 17-30% development). All fluorescent images were taken at medial wing region (M_3_/Cu_1_ crossvein area) for consistency. Scale bars= 10 μm.

Marker genes for each cluster were selected based on enriched expression within each cluster, and known marker genes during SOP specification (Figure 3C; Table S1). The identities of clusters for wing margin and trachea could be inferred based on known expression profiles. The cluster *margin* contains the genes *fz3*, *Wnt6*, *wg*, and *cut*; expression of all these genes has previously been localised to the wing margin region in multiple species of butterfly (Banerjee et al., 2023; Dohrmann et al., 1989; Hanly et al., 2023; Macdonald et al., 2010). Additional markers of this cluster included the transmembrane transporter *CG15406*, a S1A non-peptidase homolog *CG31326*, the extracellular matrix gene *M-spondin* (*mspo*) and the transcription factors *Beadex* (*Bx*) and *hairy* (*hry*). The cluster *trachea*, consisting of 302 nuclei, is marked by the expression of several *Drosophila* tracheal marker genes, the Smad-like gene *expansion* (*exp*), *chitinase 10* (*Cht10*) and *CG42749* (Devine et al., 2005; Iordanou et al., 2014).

To confirm the identity of clusters corresponding to *epithelium* and *SOP*, we focused on top differentially expressed genes in one of the two clusters, and also examined the expression of known SOP markers from *D. melanogaster*. SOP determination involves lateral inhibition between SOP and epithelial cells, where proneural genes repress *N* expression and activates *senseless* in SOPs promotes its neurogenic fate, whereas the expression of *N* in epithelial cells suppresses the fate conversion. Our data showed similar patterns of high *N* expression in the cluster *epithelium*, and high expression of proneural genes *senseless* (*sens*), *Delta* (*Dl*) and *ASH2* in the cluster *SOP* (Figure 3C). Co-immunostain confirmed Notch protein localization to membranes of epithelial cells, and non-overlapping nuclear expression of Sens (Figure 3D). Additional factors downstream of N signalling include cell adhesion molecules (CAMs), which are specialised transmembrane proteins required for proper cell sorting and SOP differentiation. The Irre Cell Recognition Module (IRM) gene family was previously implicated in cell-cell recognition and sorting events, by direct transmembrane protein-protein interactions to facilitate regular spacing of the sensory bristles in the *Drosophila* wing margin (Fischbach et al., 2009; Zhuang et al., 2009). Our data shows high expression of IRM genes *roughest* (*rst*) and *kin of irre* (*kirre*) in the cluster *epithelium*, while *hibris* (*hbs*) and *sticks and stones* (*sns*) are highly expressed within the cluster *SOP*. In addition, a homolog of *IrreC-rst* (*IrreC/rst-like*) showed enrichment in the cluster *scale-building cell*, consistent with its expression in *B. anynana* (Prakash et al., 2024). *In situ* expression of *sns* confirms its restricted expression in SOPs (Figure 3E), providing evidence that IRM genes participate in the segregation of SOPs from surrounding epithelial cells. Similarly, *E-cadherin* (*E-cad*) and *N-cadherin* (*N-cad)* are expressed in a non-overlapping fashion that are hallmarks of differential cell adhesion process in *Drosophila* wing and eye tissues (Classen et al., 2005; Hayashi & Carthew, 2004; Schäfer et al., 2014; Togashi et al., 2024), with enrichment of *E-cad* expression in the clusters *epithelial*, *socket-building cell* and *margin*, and high expression of *N-cad* in the SOP cluster. These results suggest that Notch-driven lateral inhibition is coupled with the differential expression of CAMs to drive SOP spacing and specification within the butterfly pupal wing epithelium.

Other genes showing expression restricted to cluster *SOP* include those previously described in early scale development *Hr38* and *fz2*, and the asymmetric cell division factor *inscuteable* (*insc*; Banerjee et al., 2023; Hanly et al., 2023; Le Borgne et al., 2002; Nolo et al., 2000; Prakash et al., 2024). HCR confirmed SOP-specific expression of *fz2* and a POU gene *pdm3*, with no described roles in SOP specification (Figure 3F-G). We failed to identify expression of potential pIIb markers *pros* and *repo*, suggesting that these nuclei were not captured in our dataset, possibly due to their transient nature. Cluster *epithelium* showed high expression of *Sox102F*, which has yet to be characterised for its function or expression. These results suggest that nuclei in the SOP cluster have a distinct expression profile from those in the cluster *epithelium*.

Top markers for the cluster *socket-building cell* include the genes *Su(H)* and *Sox15*, which are key transcription factors for the differentiation of the socket cell in the *Drosophila* mechanosensory bristle (Miller et al., 2009). Additional marker genes in sockets include *neuromusculin* (*nrm*), previously identified as a marker of socket cells in developing *Drosophila* tarsi, the gene *tau* for microtubule-associated protein, as well as *cypher* (*cyr*) and *unc-5* (Hopkins et al., 2023). Asymmetric cytoplasmic factors within the SOP lineage include *numb,* which is known to localise asymmetrically to pIIb and socket-building cells (Guo et al., 1996; Rhyu et al., 1994; Wirtz-Peitz et al., 2008). Socket-building cells express *N* as well, and we confirmed HCR expression of *N* in socket-building cells and epithelial cells at this stage (Figure 3K). We identified a high expression of *numb* in the cluster *socket-building cell* as well, adding confidence to the identity of this cluster.

Likewise, the cluster *scale-building cell* presents top markers including *shaven* (*sv*) and *spineless* (*ss*), two canonical markers of the shaft. Gene expression by hybridization chain reaction (HCR) revealed the concomitant expression of *sv* in the larger scale-building cells and *Su(H)* expression in smaller socket-building cells (Figure 3H). Expression of *ss* was restricted to scale-building cells similar to *sv* (Figure 3I). Within the same cluster *scale-building cell*, additional marker genes include *CG14395*, part of the gene regulatory network (GRN) including *shavenbaby/ovo*; the gene *shavenoid* (*sha*), also found in Hopkins *et al*. (2023). Cluster *scale-building cell* also expressed genes for transcription factors *ichor* (*ich*) and *couch potato* (*cpo*), the latter which was also identified in Prakash *et al*. (2024) and we confirmed its expression in scale-building cells by HCR (Figure 3J). Cytoskeletal gene *quail* (*qua*) and the lncRNA *ivory*, responsible for the normal development of scale colour, were identified as well (Hopkins et al., 2023; Kittelmann et al., 2018; Livraghi et al., 2024; Prakash et al., 2024). *Osiris4* (*Osi4*), a member of the insect-specific *Osiris* family of genes recently found to be expressed in a variety of *Drosophila* macrochaetae, was also restricted to the scale-building cell cluster (Sun et al., 2024). Altogether, expression of these genes provided confidence in the assignment of this cluster to scale-building cells.

### The specification, differentiation and development of the SOP lineage

Having assigned cluster identity to the full data set, we focused on nuclei identified as part of the SOP lineage to recover the temporal dynamics and further clustering within these nuclei (Figure 4). After subsetting the overall data for SOP, scale- and socket-building nuclei, 2,164 nuclei were retained. Performing normalisation and clustering on the subset dataset produced fourteen subclusters with marker genes exhibiting subcluster-specific expression (Table S2). We used scVelo on the newly defined subclusters, which makes use of gene splicing dynamics to predict the transcriptional state of each cell. Top genes driving the velocity prediction included *sv*, *gl-like* and *nrm* that exhibited inductive transcriptional states especially in subcluster *d* (Figure S4B-C, Table S3). After confirming that RNA velocity predicted from transcriptional states corresponded well with the progression of true time (Figure 4B), we verified that PC1 defining the subclusters corresponded to known sampling time (Figure 4C). The strong temporal component differentiating these subclusters allowed us to assign the nuclei into a pseudo-temporal series, at a higher resolution than the sampled timepoints. We recovered a ‘bifurcating tree’ model from the data with *SOP* as the initial node and *scale-* and *socket-building cell* as two terminal nodes, each corresponded by the proportion of nuclei in G1, G2/M or S phases (Figure 4C).

**Figure 4.**
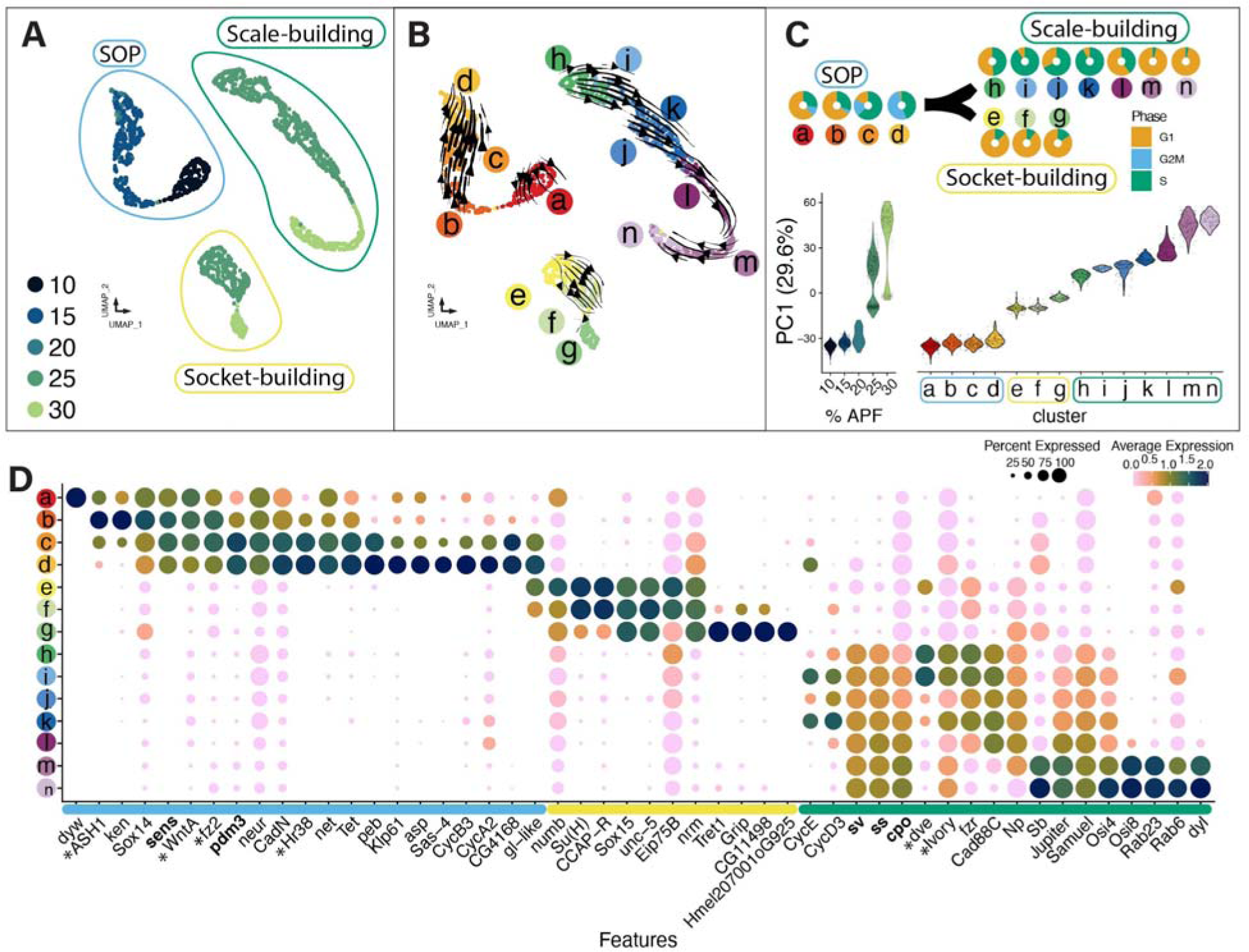
Pseudotime prediction corroborates the temporal sampling of cells identified in the SOP lineage. **A.** UMAP visualisation of nuclei belonging to the SOP lineage coloured according to sampled timepoints (10-30% pupal development). **B.** RNA velocity prediction of pseudotime ordering of nuclei within SOP, scale- and socket-building nuclei. Unsupervised clustering of SOP lineage resulted in 14 subclusters (a-n) **C.** Comparison of G1 interphase, G2M mitotic, and DNA replication S phase show cell cycle dynamics across the 14 subclusters. PC1 (29.6%) used for UMAP clustering identified a strong temporal component separating the subclusters. **D.** Each subcluster shows semi-independent gene expression profiles with non-overlapping marker genes. Genes in bold were tested for *in situ* expression in Figure 3 and genes with * have known expression data either from this paper or from previous publications (Ficarrotta et al., 2022; Hanly et al., 2023; Livraghi et al., 2024; Pomerantz, 2021; Prakash et al., 2024).

Given that Seurat-assigned subclusters tracked the progression of real time-anchored pseudotime, we explored whether cell cycle states contributed to the delineation of subclusters. Using cell cycle markers from *D. melanogaster*, we showed that subcluster ***d***, the latest time point in the SOP cluster, comprised cells either in S- or G2M-phase, and none in G1 interphase. The cell cycle state exhibited by subcluster ***d*** presents these cells as intermediate between SOP and scale- or socket-building cells, corresponding to the mitotic event during Division I. Within subclusters ***e*** to ***g*** that formed the original cluster *socket-building cell*, most nuclei appeared to be in G1, unlike those in subclusters ***h*** to ***n*** which came from the cluster *scale-building cell*. The scale-building cells exhibited an intermediate S-phase stage from ***h*** to ***k***, with none scored as entering G2M; then in the later subclusters ***l*** to ***n***, most nuclei were scored in G1. This corresponds to the massive increase of nuclear size and ploidy within the nucleus of scale-building cells, where nuclear DNA replication is not accompanied by cellular or nuclear division (Cho & Nijhout, 2013; Henke & Henke, 1946).

Additional marker genes for cluster ***d*** support this cluster as the asymmetrically-dividing SOP cells, including centriole assembly and microtubule organizing center (MTOC) genes *sas-4*, and *asp*; cytokinesis-related genes *subito* (*sub*), *fascetto* (*feo*) and *Klp61* (Figures 4D, S4). Previous work noted that G2 phase arrest was required prior to SOP fate determination (Kimura et al., 1997; Knoblich et al., 1994). We found high expression of cyclin genes *CycB, CycA2* and *CycB3* in the same subcluster ***d***, providing further support that SOP nuclei enter M phase preceding the bifurcation into the clusters *scale-* and *socket-building*. Non mitotic-related genes were expressed simultaneously in the same cluster, such as methylation factor *Tet* and transcription factors *peb*, *Hr38*, *net*, *sens*, and a homolog of *Drosophila glass* (*gl-like*), highlighting more SOP-specific markers (Figures 4, S3).

To differentiate the clusters containing *socket-* versus *scale-building cells*, we first used several markers of socket-building cells, including the canonical *Sox15 and Su*(*H*), and others *CCAP-R*, *unc-5, Eip75B* and *nrm* to confirm the identity of three subclusters ***e*** to ***g***, which originated from the socket-building cell cluster (Figures 4D, S4; Hopkins et al., 2023). Subcluster ***g*** with the most mature sockets had marker genes *Glutamate receptor interacting protein* (*Grip*)*, CG11498* and a gene with unknown fly orthologs, suggesting novel gene functions within socket-building cells. Within subclusters ***i*** to ***k***, we identified the expression of cyclin genes *CycE* and *CycD3*, both being key factors in determining the transition from G1 to S, which supports the endocycling state of scale-building cells in these subclusters (Audibert et al., 2005; Lilly & Spradling, 1996; Richardson et al., 1995; Sherr, 1994). Subclusters ***m*** and ***n*** arranged in later pseudotime appear to be enriched with genes for microtubule-associated proteins *Jupiter, Samuel, Notopleural* (*Np*), *Stubble* (*Sb*), Zona Pellucida (ZP) domain protein *dusky-like* (*dyl*) and Rab-GTPases *Rab23* and *Rab6* (Figures 4D, S3). The gene expression signatures suggest a high rate of endosomal cycling and microtubule-related processes related to scale-building within these subclusters (Adler et al., 2013; Dinwiddie et al., 2014; Tilney et al., 2000). On top of known cell fate specifying factors, the higher resolution from reclustering and pseudo-temporal series of nuclei highlighted distinct cell cycle signatures and growth-related processes.

### Marker gene validation using perturbation experiments

To validate marker genes used for cluster identification, we performed functional perturbations using CRISPR/Cas9 site-mediated mutagenesis for genes involved in the specification and differentiation of *D. melanogaster* SOP, in *P. interpunctella* and the painted lady butterfly *V. cardui*. First, the known marker gene for scale-building cells, *sv*, was knocked out using CRISPR-Cas9 mutagenesis in *P. interpunctella*. CRISPR mosaic knockout (mKO) of *sv* (n=463) was highly lethal and yielded two surviving adults. Both had scale-absent patches but normal sockets on the wings (Figure 5A-B’), replicating the results for *sv* knockout in the butterfly *B. anynana* (Prakash et al., 2024). These results confirmed the conserved necessity of *shaven* in fly bristle and lepidopteran scale formation, independent of socket formation.

**Figure 5.**
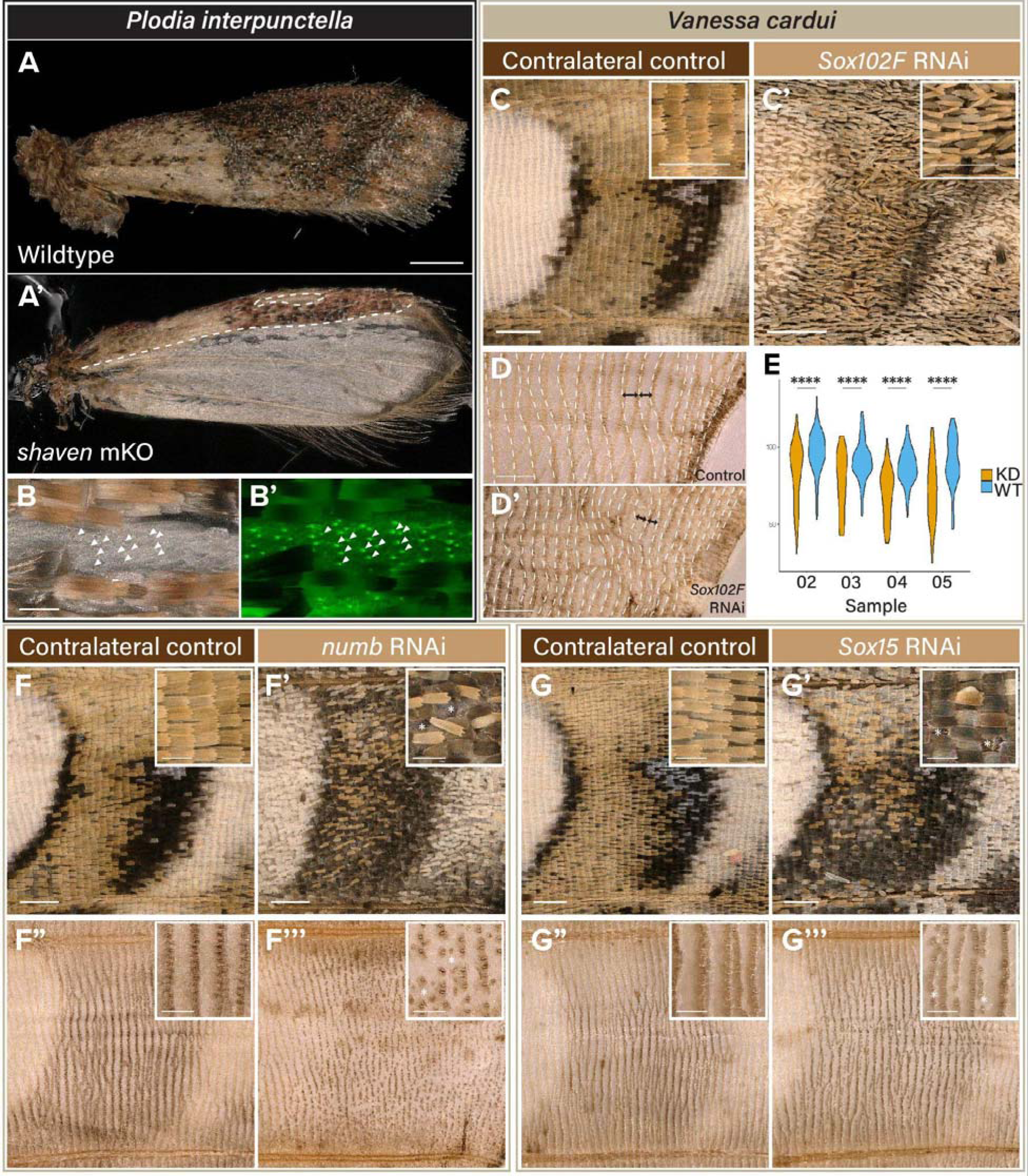
Functional perturbation of marker genes known in canonical SOP differentiation pathway resulted in scale- and socket-specific phenotypes in *P. interpunctella* (A-B’) and *V. cardui* (C-G’’’). **A-B’** Mosaic knockouts of *sv* resulted in two individuals with missing scales.Scale-deficient clones maintained proper socket formation, evident from socket green autofluorescence in B’ (triangles: examples of empty sockets). **C-D.** RNAi electroporated individuals for *Sox102F* exhibited reduced and narrowed scales that resemble underdeveloped scales (C-C’) and wings with chemically removed scales showed reduced row spacing between sockets, indicating a loss of epithelial area (double-headed arrows) **E.** Measurements of socket row spacing in chemically descaled *Sox15* RNAi knock-down (KD) wings (N = 4) compared to their contralateral wild-type controls (WT-CL). ****: Wilcoxon test, *p-value* < 0.0001. **F-F’’’** RNAi electroporation of *numb* resulted in a disorganised array and loss of scales and sockets. **G-G’’’** RNAi electroporation of *Sox15* led to loss of scales and sockets limited to the veins and specifically the loss of alternating cover scale/sockets. * indicates loss of scale and socket at the position when compared to contralateral control. All wing regions displayed for *V. cardui* were taken from the M_2_/M_3_ wing vein compartment. Scale bars: A = 500 μm; C-G” = 200 μm, C-G” insets = 100 μm.

Due to high embryonic lethality and possible pleiotropic effects of gene knockouts, we employed Dicer-substrate small interfering RNAs (dsiRNAs) to perform expression knockdown by RNA interference (RNAi) of putative marker genes for *epithelium*, *socket-* and *scale-building cell* clusters respectively, *Sox102F*, *Sox15* and *numb* (Figures 4D, 5C-F). Our snRNAseq identified high *Sox102F* expression in epithelium and later at lower levels in the SOP lineage (Figure 4D). To test a possible role of *Sox102F* in epithelial development, we used wing RNAi electroporation to knockdown *Sox102F*. All treated wings (6/6) showed narrowed and shortened scales without affecting the colour pattern or scale length (Figure 5C-C’) and overall curling unto the ventral surface that was perturbed, indicative of a reduction in wing surface due to *Sox102F* knockdown. To understand the factor causing wing surface area reduction, we chemically removed all scales to observe socket spacing that mark the position of each accompanying scale, and identified a significant reduction in spacing between 80-100 scale/socket rows in the same wing vein compartment (M_2_/M_3_), consistent in four experimented wings when compared to their contralateral controls (Figures 5D-E, S5-6). These results suggested that *Sox102F* knockdown resulted in fewer or smaller epithelial cells within the wing bilayer, but definitive evidence will require live tracking of epithelial cell development.

Progression through SOP lineage involves asymmetric division, which occurs via the polarisation of cytoplasmic components within the cell during mitotic divisions. One of the genes responsible for asymmetric division is *numb*, a membrane-associated inhibitor of Notch signalling that localises to the posterior surface of the dividing cell in *D. melanogaster* (Guo et al., 1996). Knockdown of *numb* results in transformation of all SOP daughter cells into socket cells (Rhyu et al., 1994). In *V. cardui*, knockdown of *numb* caused the disorganisation of scales across the affected wing region, complete loss of scales and sockets along the wing veins (Figures 5F-F’’’, S7). Of note, scale disorganisation was mild as the unaligned scales still pointed towards the wing margin, possibly a lower penetrant effect with the incomplete knockdown effects. Similar scale disorganisation was observed in *Sox15* knockdown wings, where perturbed areas consistently presented loss of scales and sockets on wing veins, and the loss of the cover scales and sockets in the wing, though not of ground scales (Figures 5G-G’’’, S8). This phenotype would be consistent with the failure of socket cells to properly differentiate, similar to the *Sox15^4AA^* mutant in *D. melanogaster* (Miller et al., 2009). Altogether, RNAi-mediated knockdown of marker genes revealed considerable insights into epithelial, scale and socket development in the lepidopteran wing.

### *pdm3* transcription regulates patterning and scale identity

The results outlined above emphasise that SOP specification and terminal differentiation into scale and bristle cells involve highly conserved processes between Lepidoptera and *Drosophila*. However, we also observed the expression of genes that had not been previously described in the canonical lineage, including *pdm3*. We selected *pdm3* based on its SOP-specific expression as a functional candidate to test in butterfly wing SOPs. In *Drosophila*, *pdm3* regulation was linked to the patterning of abdominal pigmentation, but with no reported effect in bristles (Rogers et al., 2014; Yassin et al., 2016). HCR profiling of *pdm3* expression at 12-14% pupal development reveals that it is spatially regulated in association with presumptive colour patterns, such as the light contours of eyespots (19-23 h APF; Figure 6A,E,F). We generated CRISPR somatic knock-outs of *pdm3* in *V. cardui* and obtained 27 surviving adults, from which 18 individuals had wing phenotypes. While we did not observe embryonic or larval lethality associated with the injections, the treated individuals showed a low emergence rate of 39% at the pupal stage, with many of the adults failing to eclose from their pupal case without manual intervention. Interestingly, all 18 crispants that successfully emerged were females, suggesting that the lethality of *pdm3* knockouts at the pupal stage was male-biased. The G_0_ mKO crispants exhibited significant darkening of patterns (Figures 6, S9), most pronounced on dorsal surfaces where orange patterns converted into a dark-melanic state, similar to the effects of *optix* gene knock-outs in this species (Thulluru et al., 2022; Zhang et al., 2017). The ventral surface also showed darkening phenotypes, but effects varied across pattern elements, with partial melanization effects that appeared most visible in the forewing ventral orange-pink pattern, as well as in the beige and ochre areas found around eyespots or in the central hindwing (Figure 6E-F). Lastly, wing marginal patterns (*i.e.* situated between the eyespot and the distal edge) were disorganised in *pdm3* crispants, with a blurring of pattern boundaries in chevron-shaped elements, and an overall expansion of blue and white fields (Figure 6C, E-F). These data implicate a transcription factor with no previous known role in butterfly wing colour patterning, and illustrate the potential of snRNAseq data in identifying important regulators of this developmental system. Future work will be required to place *pdm3* in the context of gene regulatory networks that regulate colour patterning.

**Figure 6.**
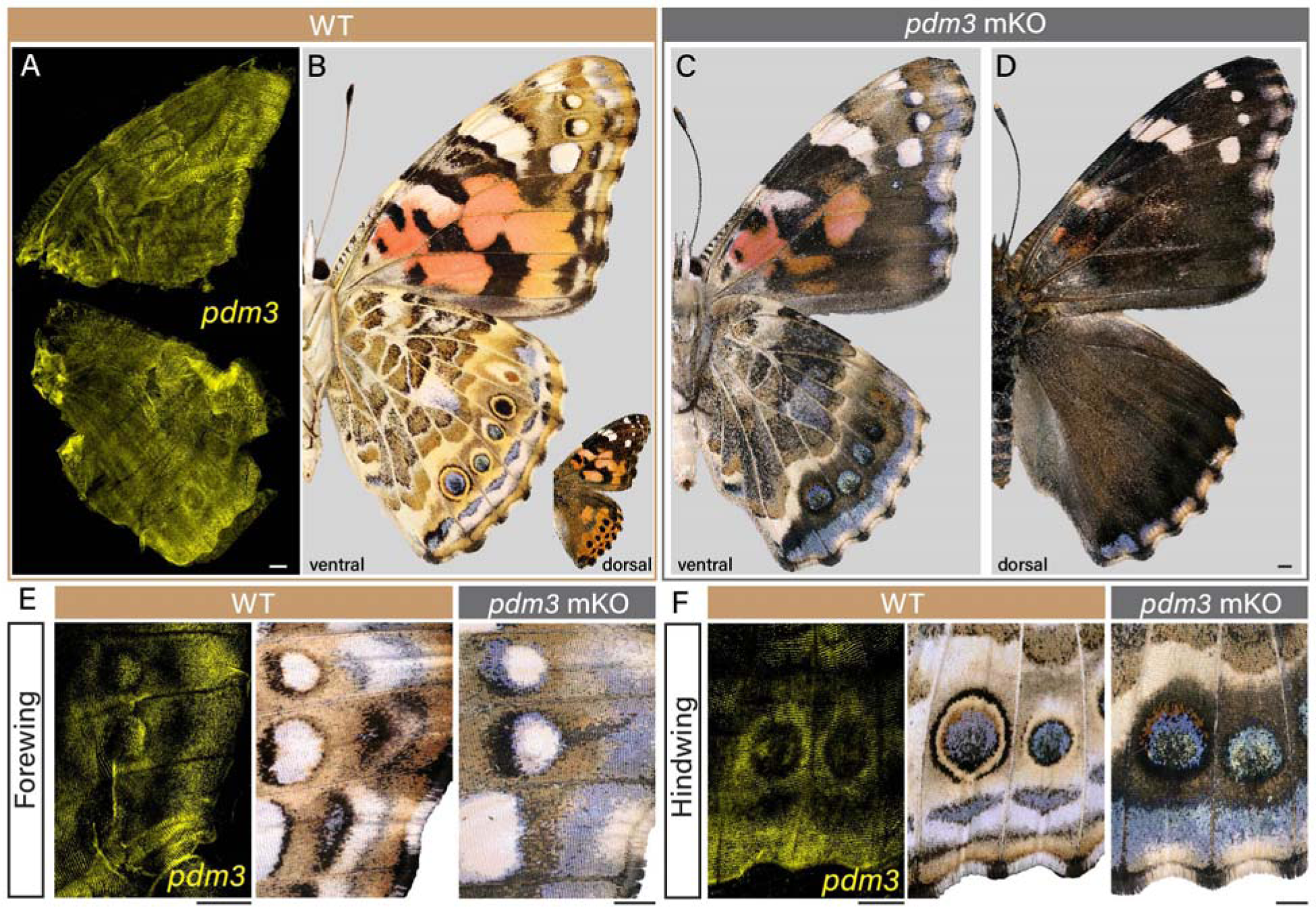
CRISPR/Cas9-mediated mKO of *pdm3* in *V. cardui*. **A-B.** HCR staining of *pdm3* expression in 12-14% pupal wings prefigures the position of adult wing pattern elements in wild type *V. cardui*. Expression is most pronounced in ventral hindwing eyespot contours, forewing eyespot white, discrete white or orange sections of the forewing, and in the marginal areas. **C-D.** Dorsal (C) and Ventral (D) views of *pdm3* mKO reveals a widespread gain of dark melanic scales on both surfaces, as well as a disruption of ventral pattern element boundaries. **E-F.** Insets of a ventral forewing (E) and hindwing (F) showing *pdm3* expression using HCR and both adult wildtype and *pdm3* mKO comparisons. Scale bars = 500 μm.

## Discussion

### Ontogeny and single-cell resolved gene expression of lepidopteran wing scales

Here, we were able to recover the ontogeny and developmental trajectory of the SOP lineage from early specification through the differentiation of the scale and socket. Our data allow us to couple differentiation events to changes in gene expression over time, providing supporting genetic evidence for homology with the *Drosophila* canonical SOP lineage, and uncovering novel effectors of pattern and colour. Our evidence broadly supports a previously described model for SOP divisions as proposed in Stossberg (1938) and recapitulated in McDougal *et al*. (2021). This model includes a division and apoptosis of an earlier cell that resembles the pIIb cell, followed by later division of the pIIa cell into a socket-building cell and scale-building cell.

Live imaging using Hoechst33342 staining in both *T. ni* and *J. coenia* pupae highlighted specification and asymmetric divisions of the SOP to produce a pIIa cell, and in *T. ni* we observed a transient pIIb cell that appeared to undergo apoptosis. We infer that the production of a pIIb is necessary for the asymmetric localization of cytoplasmic determinants that must be excluded from the pIIa cytoplasm, such as cortical cell fate determinants numb and prospero, for correct differentiation and division into scale- and socket-building cells. However, we did not directly observe the existence of a pIIb cell during the live imaging of *J. coenia*, or in HCR imaging in *V. cardui*, possibly due to the perpendicular plane of division that could make it difficult to observe a pIIb cell embedded within the epithelium. Moreover, in the snRNAseq data from *H. melpomene*, no nuclei could be assigned to a pIIb cluster. This might be due to the highly transient nature of pIIb not being captured in the time series data. Alternatively, the putative pIIb could be anucleate, which would not be captured in a single nucleus RNAseq experiment. A putative pIIb may therefore be observable with higher resolution time series using membrane markers during the rapid SOP differentiation stages in butterflies, but this does indicate a potential spatial or temporal difference with moths *T. ni* and *E. kuehniella*, where the putative pIIb can be readily observed (Stossberg, 1938). Despite this, this body of evidence provides strong support for the model in Fig 1b, where the lepidopteran scale organ is produced by a stereotypical set of ontogenetic processes including apoptosis of the pIIb.

### SOP determination and division involves canonical players of cell cycle transition and endocycling

By inferring a cell cycle phase and a pseudotime state for each cell in the SOP lineage, we observed the timing of mitotic division and the endocycling phase of nuclear genomic replication without cytokinesis (Edgar et al., 2014; Zielke et al., 2013). Similar to bristles and sockets in *Drosophila*, lepidopteran scales and sockets undergo endoreplication to produce enlarged polyploid nuclei (Audibert et al., 2005; Cho & Nijhout, 2013; Edgar & Orr-Weaver, 2001; Hartenstein & Posakony, 1989). We observed evidence of endocycling in lepidopteran scales, via the skipping of G2/M by cycling between G1 and S phase. This event begins around the time of formation of scale-building cells and appears to cease in some subclusters at 30%, which then exits into a non-cycling G1 phase, matching previous observations of endoreplication in the scale cell (Cho & Nijhout, 2013). In *Drosophila*, specific timing of mitotic divisions is critical to set up the final positions of adult bristles, the latter which are fixed and important for proper axon pathfinding from the brain to the sensory organ (Smith & Sondhi, 1961). Understanding the combinatorial effects of cellular divisions and endoreplication will provide further mechanistic insights into how lepidopteran scale-building cells are derived from a mother SOP.

### The Scale cell lineage displays a common gene expression cascade with other sensory organs

The combination of lineage reconstruction and single nucleus transcriptomics provided temporal anchoring of key events during scale organ development, highlighting the relative timings of mitotic divisions to the expression of marker gene candidates. For early SOP specification, many genes identified herein recapitulated gene expression common to those in *Drosophila* mechanosensory bristles, like the recruitment of Notch signalling in cis-inhibition. Additionally, we found evidence for the involvement of differential cell adhesion during SOP specification from the epithelium. Indeed, cell adhesion factors like the IRM proteins and cadherins showed cell-type specific expression profiles that are consistent with their known roles in epithelial-SOP cell sorting during bristle development (Linneweber et al., 2015; Takemura & Adachi-Yamada, 2011). We also recovered the later division of SOP-II into the scale-building ‘trichogen’ and socket-building ‘tormogen’ cells, and detected expression of genes with described roles in bristle development, including insect-specific *Osi* genes, microtubule-associated genes and endosomal sorting components including Rab-GTPases. Altogether, both data corroborate key cellular events with distinct gene expression signatures at specific time periods during wing development.

Beyond expression profiles, functional experiments validated the roles of key marker genes, recovering scale-loss when knocking out *sv* and loss-of-socket effects when knocking down *Sox15.* However, the incomplete knockdown effects of *Sox15* in removing sockets may mean that other factors are required to abrogate socket cell formation, as observed in *Drosophila* where socket cells still formed in *Sox15* mutant lines (Miller et al., 2009). Functional experiments of other genes in *Drosophila* bristles suggest that both intrinsic and extrinsic factors are involved in cell fate decision making (Heitzler & Simpson, 1991; Posakony, 1994). Intrinsic signals are specified in one daughter cell but not the other, like genes involved in asymmetric partitioning of genes during mitotic division (Rebeiz et al., 2011), while extrinsic signals are acquired via cell-cell signalling (Heitzler & Simpson, 1991). Our validation results of some of the known intrinsic and extrinsic signals suggest that different types of factors facilitate fate commitment to different degrees.

In sum, SOP specification and division involves the rapid and robust processes, which emerge from the coordination of rapid cellular division or endocycling states via oscillating cyclin/CDK signals, correct asymmetric localization of cytoplasmic components, cell-cell signalling, and proper cell-cell contacts maintained by CAMs. We observed both intrinsic, autonomous factors and extrinsic, signalling factors during lepidopteran scale development from the gene expression profiles and functional data, thus provide corroborating evidence that the lepidopteran scale is part of the canonical lineage of sensory organs as defined by Lai & Orgogozo (2004).

### Sockets, Scales, and the evolution of sensory organs

SOP derivatives like scales, setae and sensilla are quintessential examples of serial homology, a notion that captures the shared origin of repeated but specialised anatomical parts (Wagner, 2014). Serial homologues and cell types can be treated as evolutionary characters and degrees of developmental relatedness that can be traced on a phylogenetic tree (Arendt et al., 2016; DiFrisco et al., 2023). This framework notably extends the notion of orthology and paralogy from genes, to serially homologous biological processes such as lineages and cell types. For example, we can now affirm that socket precursor cells are cell type “orthologs” between Lepidoptera and Diptera, not only because of their conserved placement in the SOP lineage, but also because they share the markers *Su(H)* and *nrm,* regardless of whether they are associated to scales (this study), or sensilla and combs (Hopkins et al., 2023). Meanwhile, the shaft component of *Drosophila* sensilla (whether mechano- or chemosensory) and the non-sensory comb can all be seen as “paralogs”, co-existing within a lineage as specialised derivations of the SOP canonical archetype.

We can extend this reasoning to a given SOP cell and its descendants – including their neuronal, glial, scale and socket cell outputs – meaning that specialised sensory organs and other SOP-derived serial homologues can be treated as paralogous within a species. From a macroevolutionary perspective, our data validates the serial homology of lepidopteran scales with dipteran sensory bristles. However, it is unclear if these represent a case of orthology, because scales may have instead evolved from a non-sensory serial homologue that pre-existed in a common ancestor, akin to the ornamental setae and scales found in bumblebees and mosquitoes (Djokic et al., 2020; Hines et al., 2022).

Within Lepidoptera, we propose that the prolific diversification of scale shape and colour phenotypes boils down, in essence, to a diversification of cell type paralogs. The single-nucleus approach successfully allowed the isolation and profiling of scale precursor cells across successive stages, an achievement that may not be possible with the dissociation methods using single-cell approaches. We envision that single-nucleus transcriptomics, performed across lepidopteran species, will decipher scale subtypes with evolutionarily traceable identities, notably by enabling the identification of regulatory factors that are required for their specification and phenotypic divergence. Amidst an increasing interest in using these new technologies to build phylogenetic trees of eukaryotic cell types (Church et al., 2024; Tanay & Sebé-Pedrós, 2021), the simplicity and diversity of the SOP canonical lineage could thus provide a powerful comparative framework for evolutionary cell biology.

### Early competence in the undifferentiated SOP modulates scale colour fate

Of note, the snRNAseq dataset revealed novel insights into the development of butterfly wing scales. We identified pattern-related genes restricted to the early SOP cells like *WntA* and *fz2*, and *Ivory* in the later scale-building cells, the latter specifically implicated in melanic patterns across Lepidoptera (Banerjee et al., 2023; Fandino et al., 2024; Hanly et al., 2023; Livraghi et al., 2024; Tian et al., 2024). Additionally, we identified a gene, *pdm3,* that contributes to scale cell development and differentiation, which is not found in analogous contexts in *Drosophila.* Instead, *pdm3* was implicated as a repressor of dark pigmentation across several *Drosophila* lineages (Rogers et al., 2014; Yassin et al., 2016). In our case, *pdm3* expression is restricted to the earlier SOP cell and was not detected at later stages in differentiated scale-building cells. Knocking out *pdm3* in *V. cardui* led to darkened colour and alterations to specific patterns, without disrupting proper scale formation, suggesting that *pdm3* may function as a repressor of dark melanic pigments as in *Drosophila*. Given that a variety of factors like *pdm3*, *WntA* and *fz2* are expressed at this early pre-division II stage and downregulated by the time the scale begins to develop, fates of scale-building cells appear to be derived when SOPs adopt intermediate cellular states and divide. As such, there must be factors that persist through endocycles to maintain this positional identity to the scale-building cell on the wing. The plasticity of the SOP lineage in producing diverse sensory organ types appear to stem from the cellular events occurring concurrently with the asymmetric division of the SOP and later endocycling scale-building cell. Illuminating the spatial and lineage relationships between scale cells through development will provide further insights into how specification and differentiation programs evolve to generate diversity of form and function.

## Methods

### Animals

Individuals from *Heliconius melpomene rosina* (Boisduval) and *H. m. ecuadorensis* (Emsley) were reared in the insectary facilities in Smithsonian Tropical Research Institute between March and April 2022. Larvae were fed on their preferred host plant *Passiflora morifolia*. Pupae were transferred to incubators set at 28°C, 80% relative humidity, 12:12 h light cycle, in which total developmental time was measured to 200 h (∼8.5 days), from pupa formation to adult eclosion.

*Vanessa cardui* (Linnaeus) butterflies were purchased from Carolina Biological Supplies and reared at 25°C with a 16:8 h light cycle on a multiple-species diet (Southland Products Inc.). *Plodia interpunctella* (Hübner) moths were reared at 28°C as previously described (Heryanto et al., 2022; Heryanto et al., 2022). *Trichoplusia ni* (Hübner) were ordered from Frontier Agricultural Sciences (L9282) and reared on the supplied diet until pupation. *Junonia coenia* (Hübner) originated from the laboratory colony of Fred Nijhout (Duke University) and were maintained on a multiple-species diet (Southland Products Inc.) mixed with powder made from dried *Plantago lanceolata* leaves.

### Time-lapse microscopy

Freshly eclosed pupae of *T*. *ni* and *J*. *coenia* were prepared for live imaging as described in (Ohno & Otaki 2015) with the modification that 0.1 mg/mL Hoechst33342 in 70% DMSO:30% Grace’s Insect Media (Gibco) was injected through the hindwing peripodial membrane using a pulled glass capillary needle in lieu of soaking. Injected pupae were placed into glass bottom dishes and covered with damp cotton to maintain humidity during imaging. Two-photon imaging of nuclear divisions began within 45 min of pupa formation at the earliest and was conducted on a ZEISS LSM710 inverted confocal with Coherent Chameleon Vision II laser, 20X 0.8 M27 objective, and Non-Descanned Detector consisting of a two-channel reflected light GaAsP detector. A laser wavelength of 722 nm was used for two-photon excitation of Hoechst33342 dye at ∼2-4% laser power.

### Frozen nuclei extractions

For tissue collection, both forewings were dissected from male pupae sampled between 10% and 30%, for a total of 7 samples at 5% developmental intervals (10 h intervals at 28°C), and flash-frozen in liquid nitrogen before storage at -80°C. Nuclei isolation protocol was adapted from McLaughlin *et al*. (2021). Briefly, flash-frozen wings were thawed on ice for 10 min, supplemented with 200-500 μL homogenization buffer (250 mM sucrose, 10 μM Tris pH 8.0, 25 mM KCl, 5 mM MgCl_2_, 0.1% Triton-X 100, 40 U/μL RNasin Plus, 1x Protease Inhibitor, 0.1 mM Dithiothreitol), and mechanically lysed using loose (A) and tight (B) dounce homogenisers until a cloudy suspension was observed. Isolated nuclei were spun down at 1,000 x g in a 4°C centrifuge for 5 min, upon which the supernatant was removed and replaced with a nuclei suspension buffer (NSB; 1X PBS, 1% Bovine Serum Albumin, 40 U/μL RNasin Plus) before filtration through a PluriStrainer^®^ 40-μm mesh.

For samples at 25% and 30% pupal development, nuclei suspensions in NSB were not filtered, and instead centrifuged on a sucrose gradient to remove cellular debris including scales (Sigma Aldrich, Nuclei Pure Prep Nuclei Isolation Kit, NUC-201). Following manufacturer’s recommendation, 500 μL nuclei suspension was mixed with 900 μL of 1.8 M sucrose solution, before layering onto 500 μL of 1.8 M sucrose solution. The sucrose gradient was centrifuged at 4,500 × g for 45 min at 4°C on a slow acceleration set-up and without deceleration brakes. Supernatant containing the cellular debris was removed, leaving ∼40 μL, before resuspending the nuclei pellet in 500 μL NSB and transferring the solution to a 1.5 mL LoBind tube. Resuspended nuclei were centrifuged at 500 × g for 5 min at 4°C, after which the pellet was resuspended in 100–300 μL NSB and filtered through a PluriStrainer^®^ 40-μm mesh. 10 μL aliquots were drawn from each nuclei suspension to stain with Trypan Blue before manual counting with a hemocytometer. Nuclei suspensions were diluted to between 700 and 1,800 nuclei/μL and a library preparation was performed with a targeted recovery of 5,000 nuclei.

### Library preparation and cDNA sequencing

Library preparation was performed using the v3 Chromium Single Cell 3’ Reagent Kit for the 10x Genomics 3’ Gene Expression experiment. Final cDNA library concentrations were quantified using Qubit and fragment sizes were assessed with the Agilent 2100 BioAnalyzer High Sensitivity DNA kit (5067-4626). Libraries were pooled on an Illumina NovaSeq 6000 S4 flow cell. Sequencing was performed at an average sample depth of about 281 million reads and an average of 56,250 100-bp paired-end reads/nuclei by Duke University Sequencing and Genomic Technologies (SGT).

### Single-nucleus RNAseq analysis

BCL files were converted using bcl2fastq2 (RRID:SCR_015058) and aligned to the *H. melpomene melpomene* v2.5 genome (Martin et al., 2019) with the *H. melpomene melpomene* v3.1 annotation (Cicconardi et al., 2023). The annotation orthology assignment was supplemented with blastp alignment to all *Drosophila* polypeptide sequences (Camacho et al., 2009). Alignment was performed with STARSolo, counting ‘genefulls’ that permits the counting of intronic reads (Kaminow et al., 2021).

Data analysis was performed using Seurat v5 (Hao et al., 2024). First, all *STARSolo* filtered matrices were made into *Seurat* objects, filtered, normalised with *SCTransform* and merged (Figure S3). Additionally, integration was performed to account for variation between the two pattern types (*H. m rosina* and *H. m ecuadorensis*). Cell cycle scores were calculated using Seurat function *CellCycleScoring* from the expression of cell cycle markers identified in *Drosophila melanogaster* (Steinbaugh, 2018). UMAPs and k-means clusters were generated using the first 12 principal components with a resolution of 0.5. Markers were selected with FindAllMarkers, using an avg_log2FC cutoff at 0.5.

### Fluorescent in situ hybridizations

Hybridization Chain reaction (HCR) probes were designed against exonic sequences of genes, using the tool *insitu_probe_generator*, permitting a GC content in the range 35-75% and allowing limiting runs of poly-GC and poly-AT to a maximum of 3 bp (Kuehn et al., 2022). Between 6 and 10 probe pairs were designed per gene, for the genes *ss*, *pdm3*, *sv*, *cpo*, *Notch*, *sns* and *Su(H)* (Table S4). Pupal wings were dissected from the pupal case in cold 1X PBS as previously described (Hanly et al., 2023), transferred to a fixative solution (750 μL PBS 2 mM EGTA, 250 μL 37% formaldehyde) containing 9.25% formaldehyde at room temperature for 30 min, washed four times in PBS containing 0.01% Tween20 (PBT), permeabilised in 1 μg/μL of ProteinaseK diluted in PBT solution, and washed with a stop solution containing PBT and 2 mg/mL glycine and followed by two additional PBT washes. After transferring wings to a post-fix solution (850 μL PBT, 150 μL 37% formaldehyde) containing 5.55% formaldehyde for 20 min, wings were washed four times with PBT before following the rest of the protocol as in previously published procedures (Bruce et al., 2021; Choi et al., 2018).

### Immunofluorescence

Immunostainings were performed as previously described (Ficarrotta et al., 2022). All samples were incubated in DAPI diluted in 50% Glycerol/PBS (pH 7.4) for 15 min at room temperature or overnight at 4°C, prior to mounting in 70% Glycerol/PBS (pH 7.4). Confocal imaging was performed at 60X magnification under the Olympus FV3000 confocal microscope, 10X (Plan-APO 0.45; for whole-wing tiles) and 40X (Plan-APO 1.4 Oil) objectives using a Zeiss Cell Observer Spinning Disk confocal microscope (GWNIC). Antibodies used for immunofluorescence include a rat monoclonal anti-Tubulin (BioRad #MCA77G, 1:100 dilution), rabbit polyclonal anti-Senseless (gift from Perry lab; 1:400), rabbit polyclonal anti-Beta Catenin (Sigma-Aldrich #C2206, 1:100), mouse monoclonal anti-Notch (DSHB C17.9C6, 1:5).

### CRISPR somatic knock-outs

CRISPR experiments were performed in *P. interpunctella* and *V. cardui* following previously detailed procedures (Hanly et al., 2023; Heryanto, Hanly, et al., 2022). In brief, syncytial embryos were microinjected within 15 min to 3h after egg laying with a protein-sgRNA duplex of Cas9-2xNLS (UC Berkeley QB3, 500 ng/μL) and synthetic sgRNA (Synthego, 250 ng/μL). The sgRNA target sequences were 5’-GGTGGCGACACCCCCTGTGG**TGG**-3’ for *P. interpunctella sv*, and 5’-CAGCGCTTGAGGCGTGAATG**AGG**-3’ for *V. cardui pdm3* (PAM sequences in bold).

Freshly eclosed adult *P. interpunctella* and *V. cardui* were carefully handled to minimise scale-loss independent of functional perturbations. *P. interpunctella* pupae were moved to 20°C upon pigment darkening and once eclosed, adults were frozen at -30°C. For *V. cardui*, newly eclosed adults were moved to 4°C to dry for 1 day before freezing at -20°C. Wings were imaged on Keyence VHX-5000 microscope at the 50X and 200X magnification setting with VH-Z00T and VH-Z100T lenses. Socket autofluorescence was imaged in the GFP channel of a Olympus BX53 fluorescent stereoscope mounted with a UPlanFL10X objective lens and illuminated under X-Cite 120 LED Boost at full-power.

### RNAi electroporation

Electroporation was performed as per previously reported (Hanly et al., 2023). Briefly, fresh pupae (< 5 min post pupation) were moved to 4°C for 10 min to immobilise the pupae. Once immobilised, each pupa was laid with its right side up and forewing was lifted with the cuticle, then laid onto a moist agar pad (1% agarose in 10X PBS). Injection of 2 µL of 100 µM dsiRNA was performed distal to the wing margin (apoptotic cells prior to eclosion), before a 1X PBS droplet was placed on the wing. Positive electrode was placed on the PBS droplet and negative electrode on the moist agar pad to target the ventral surface, before applying 5 pulses of 12V for 280 ms, at an 100-ms interval. PBS droplet was pipetted out before putting the forewing back into the pupal case. dsiRNA sequences are listed in **Table S5**.

## Acknowledgements

We thank A. Mazo-Vargas, L. Livraghi, M. Chatterjee, C. Arias, B. Hopkins, A. Mackay-Smith, K. Pipho, and F. Cicconardi for many intellectual and technical inputs during this project; R. Mauxion, R. Canalichio, as well assistant personnel of the Gamboa Heliconius Insectaries (STRI Panama) and Harlan Greenhouse (GWU), for help rearing host plants and butterflies; the GWU Genomics Core for performing library preparation of snRNAseq samples; the Duke University Sequencing and Genomic Technologies (SGT) for performing sequencing; P. Hernandez, A. Jeremic, and the George Washington University Nanofabrication and Imaging Center (GWNIC) for providing access to confocal microscopes; the GWU HPC team for providing computational infrastructure.

## Data availability

Supplementary Tables S1-S3 and genome annotation files for this paper are available on the Open Science Framework with DOI: 10.17605/OSF.IO/CWB6R. Sequence files are uploaded to SRA under the Bioproject PRJNA1118741

## Supplementary Materials and Methods

### Post-processing of time-lapse microscopy images

Live image acquisitions were saved as .TIF hyperstacks and processed using Fiji open source software (Schindelin et al., 2012). Correct 3D drift was run to align frames and frames that experienced too much movement and did not have discernible morphology were removed from the movie file after timestamps had been added to keep a proper morphological timeline (Parslow et al., 2014). Frames below the nuclei/scale bases were manually removed to reduce noise in projected views, making sure not to clip any sections of wing membrane epithelial cells. Frames containing patches of autofluorescence from overlying cuticle, usually as a result of the mounting having been tilted, had these areas manually excised. Once a stack was trimmed, maximum intensity or standard deviation intensity projections were used to better visualise development and a 1.25σ Gaussian blur applied to aid visual particle tracking. To visually trace lineages of nuclear divisions, representative nuclei demonstrating different division regimes were manually cropped from drift-corrected movies and the focal nucleus centered using a custom landmark-based registration plugin called Manual_Registration.py that aligns features based on user-selected ROIs (Eglinger, 2018). The focal nucleus was outlined with an ROI for all timepoints in a movie with the desired LUT. This selection was then copied into a grayscale version of the same movie to leave only the desired elements in colour.

### Data analysis for RNA velocity

To run scVelo on nuclei that form the SOP lineage (Bergen et al., 2020), sample fastq files were aligned with *Cellranger* v.5.0.0 with reference genome *Hmel v2.5*, additionally with *–include-introns* to include intronic reads. Each of the output filtered UMI count matrices was used as input for *velocyto* (La Manno et al., 2018) with the parameter *run10x* to generate uniquely mapped reads from cellranger output UMI matrices, that align to both exonic and intronic regions. The output loom files were aggregated unspliced and spliced UMI count matrices for each sample. The loom files and integrated Seurat object were read into python v3.8.17, and analysis was performed based on UMAP coordinates obtained from Seurat. scVelo v0.3.0 was used. *scvelo.utils.clean_obs_names* was invoked independently for the integrated Seurat object and concatenated loom files, before using *scvelo.utils.merge* to concatenate the processed Seurat object with loom files for all samples. In total, the aggregated *anndata* matrix contained the spliced and unspliced count matrices for all samples, forming a count matrix of 2,164 cells by 14,921 genes. The *anndata* object was prepared using *scvelo.pp.moments* and *scvelo.tl.recover_dynamics*, before the actual velocity estimation using *scvelo.tl.velocity*. All models (stochastic, deterministic, dynamical) were tested before the final choice of the dynamical mode because it recapitulated the sampled timepoints accurately using top driver genes with some known marker genes (Table S3). Velocity vectors were plotted onto the UMAP embedding and latent time for each cell was estimated using *scvelo.tl.latent_time*.

### Genotyping of crispant adults

For testing the presence of on-target mutations in *pdm3* crispants, a leg was removed from the frozen adult bodies and submerged in 19.5 μL DNARelease Buffer and 0.5 μL DNARelease (Phire Animal Tissue Direct PCR Kit, Thermo Scientific), incubated at room temperature for 2 min before incubation at 98°C for 3 min. Supernatant was diluted in 5 μL of molecular grade water before 1 μL was added to a 19 μL Phire PCR reaction (30 cycles). PCR was performed using primers (*M13*-Forward: 5’-*TGTAAAACGACGGCCAGT*CTGTATCCTTCCAGGTACGC-3’, Reverse: 5’-TGGCGAATGTCCTTGGCAAT-3’. Gels with 0.5X TBE buffer and 1.5% agarose were used for confirming PCR products, before gel extraction using the Zymoclean Gel DNA Recovery Kit. Purified gel extracts were sent for Sanger sequencing (Azenta Life Sciences). Synthego ICE analysis was performed to confirm the presence of indels within sequenced read traces (https://ice.synthego.com/).

## Supplementary Material

**Movie S1:** Continuous live imaging of *T. ni* pupal hindwing from 1 h to 24 h APF (0.7%-17% development). Hoechst33342 dye was used as a live nuclear stain and a single SOP nucleus is pseudocolored in blue. A first division resulted in SOP-II and a transient pIIb that apoptose, following a second division that results in a scale- and socket-building cell.

**Movie S2:** Continuous live imaging of *J. coenia* pupal hindwing from 24 h-72 h APF (13-39% development), with SOP nucleus pseudocolored in blue. The SOP nucleus undergoes partial nuclear condensation with no evidence of a division, followed by a division that results in a scale- and socket-building cell.

**Movie S3:** Original, uncropped view of *T. ni* pupal hindwing development from which S1 was derived, spanning 0.7%-17% development.

**Movie S4:** Original, uncropped view of *J. coenia* pupal hindwing live imaging in one individual from 1 h to 40 h APF (0.5%-13% development) and another individual from 24 h to 72 h APF (13%-39% development). The first time frame (1 h to 40 h APF) shows nuclei rotating in place with no clear division, and later the arrangement into rosette-like proneural clusters. The second time frame (24 h to 72 h APF) shows division II into scale- and socket-building cells. Separate movies were obtained from different individuals due to phototoxicity during imaging of *J. coenia*, which has a longer pupal development time. Movie S2 was cropped from the second individual in this video.

**Figure S1.**
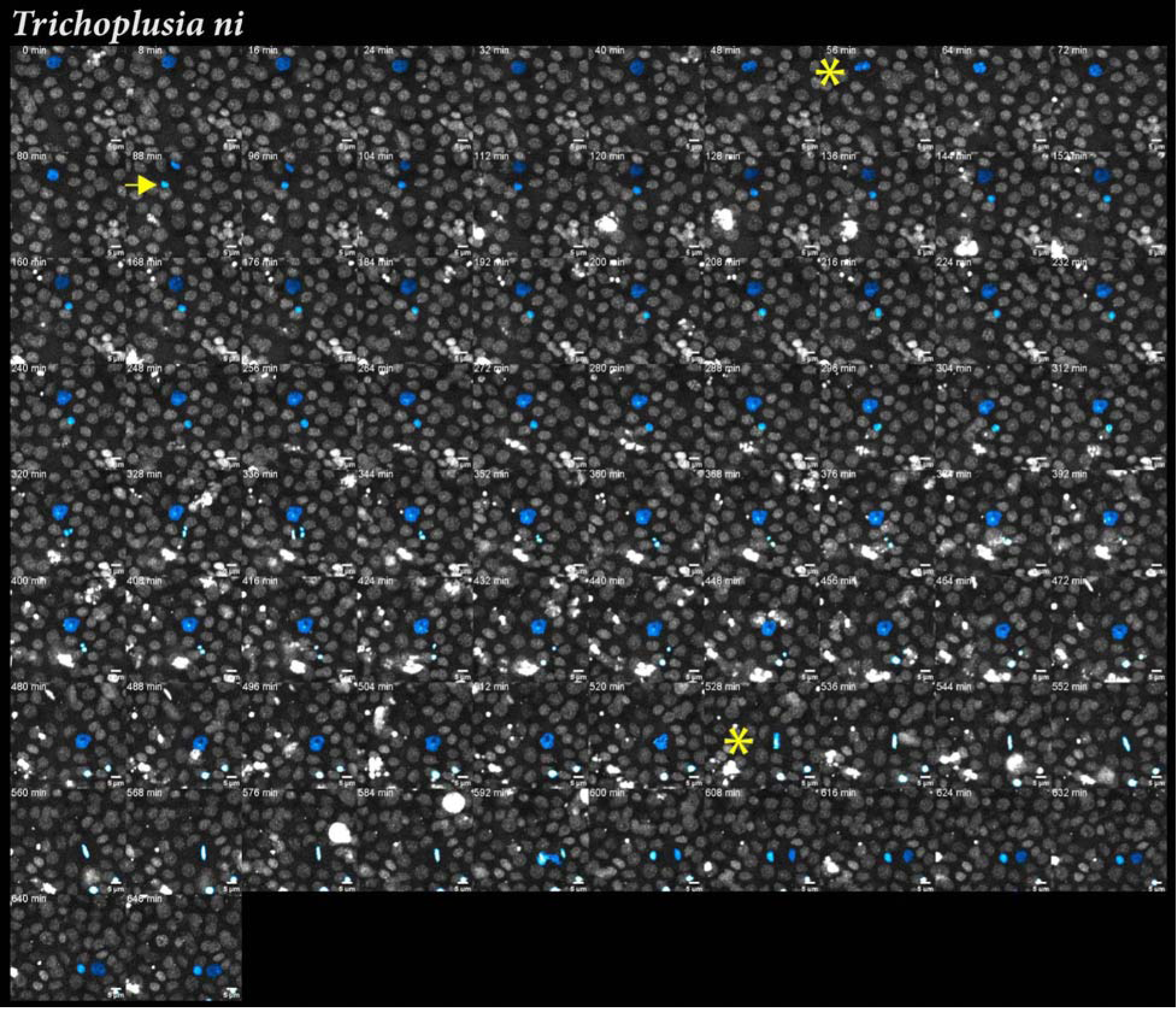
Live imaging of developing *Trichoplusia ni* pupal wing epithelium by frame. Continuous imaging of pupal hindwing from 1 h (0.7% development) to 24 h APF (17% development). Frames with asterisks denote onset of mitotic division of nucleus highlighted in blue. Arrow points to the putative pIIb. Scale bars = 5 μm.

**Figure S2.**
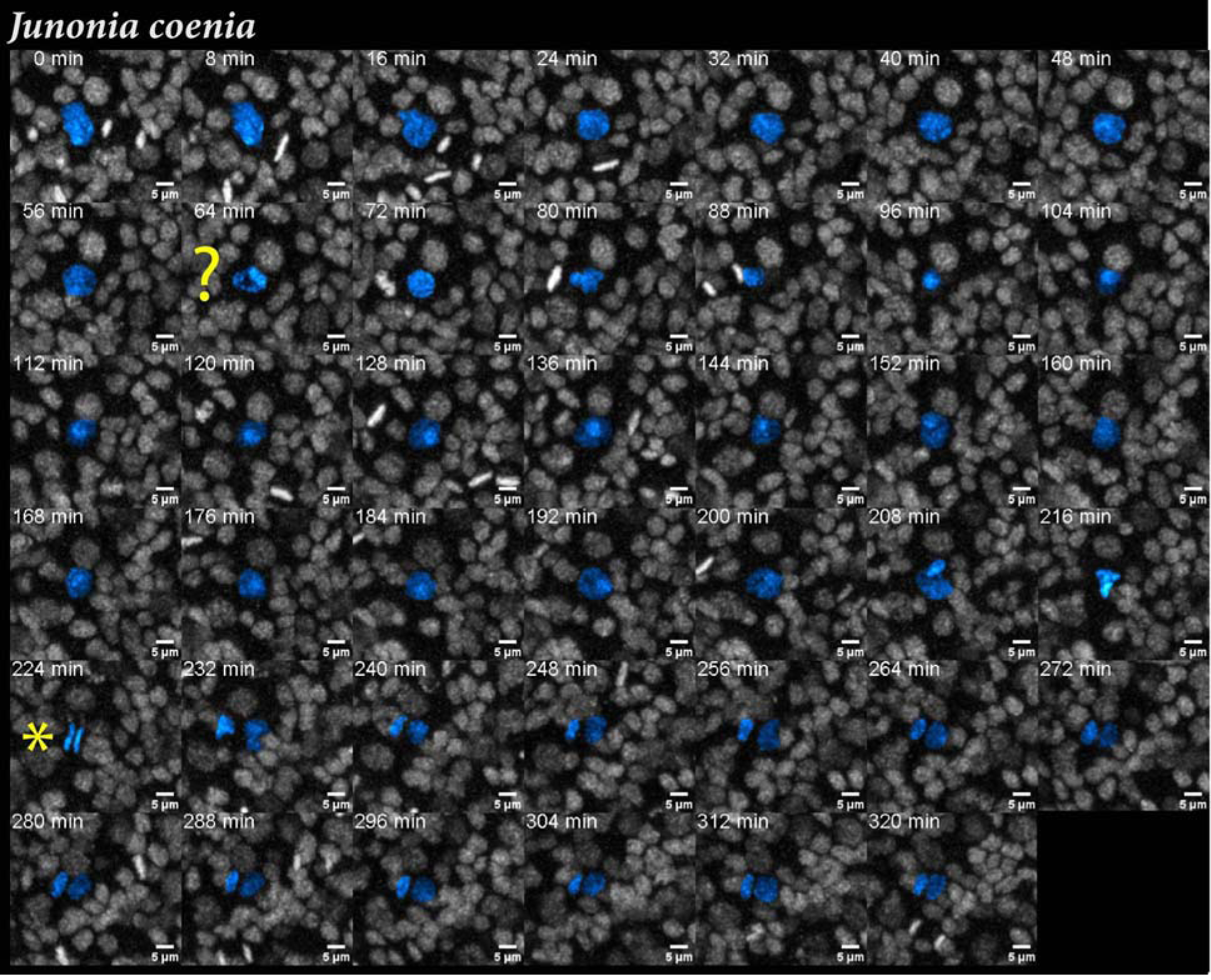
Live imaging of developing *Junonia coenia* pupal wing epithelium by frame. Continuous imaging of pupal hindwing from 24 h (13% development) to 72 h APF (38% development). Frames with asterisks denote onset of mitotic division of nucleus highlighted in blue. Question mark refers to an inconclusive mitotic division event. Scale bars = 5 μm.

**Figure S3.**
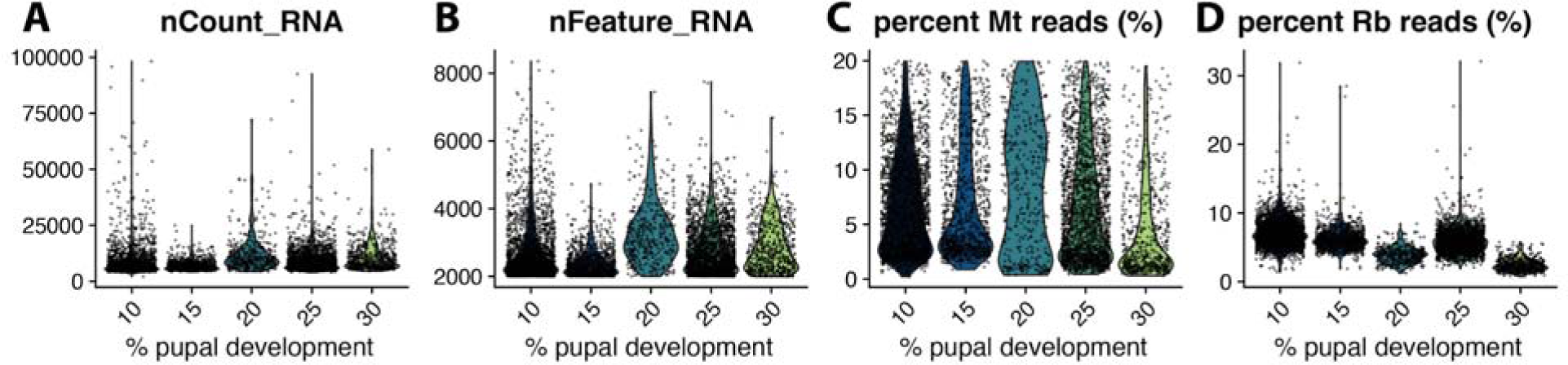
Quality control (QC) metrics for merged and integrated snRNAseq data using *H. melpomene* wing tissues. **A-B.** Genes expressed in minimally 3 cells (A) and cells with a minimum of 2000 genes expressed were retained. **C.** Nuclei with a maximal mitochondrial (Mt) read percentage 20% were retained for further analysis. **D.** Ribosomal (Rb) read percentage per nucleus was verified for each sampled timepoint.

**Figure S4.**
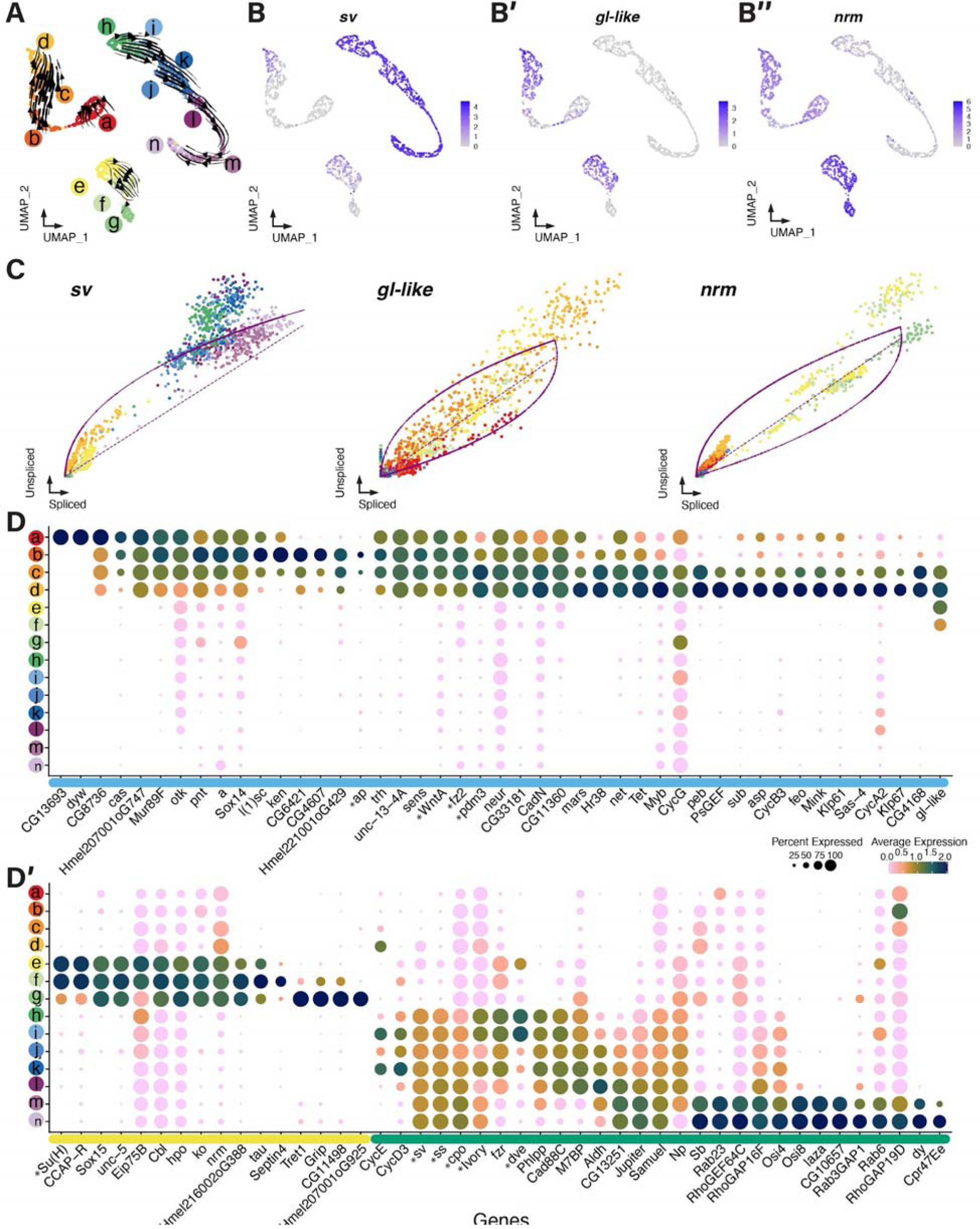
Additional genes identified in SOP lineage that track pseudotime inference using RNA velocity. **A.** Reclustering of cells forming the SOP lineage reveal subclusters that exhibit a specific pseudotime trajectory as inferred from scVelo dynamical model. **B-C.** Genes *sv*, *gl-like* and *nrm* show expression in subclusters forming the late SOP (subcluster *d*) and either scale-(*sv*) or socket-building cell (*gl-like, nrm*). Phase portraits showing unspliced/spliced counts of the three genes with a transcriptional inductive phase (unspliced>spliced) in either the *scale-building* subclusters *h-n* (*sv*) or *socket-building* subclusters *e-g* (*gl-like*, *nrm*). **D-D’.** Genes with expression specific to cluster *SOP* (D) and clusters *socket-* and *scale-building cell* (D’). (continued from previous page)

**Figure S5.**
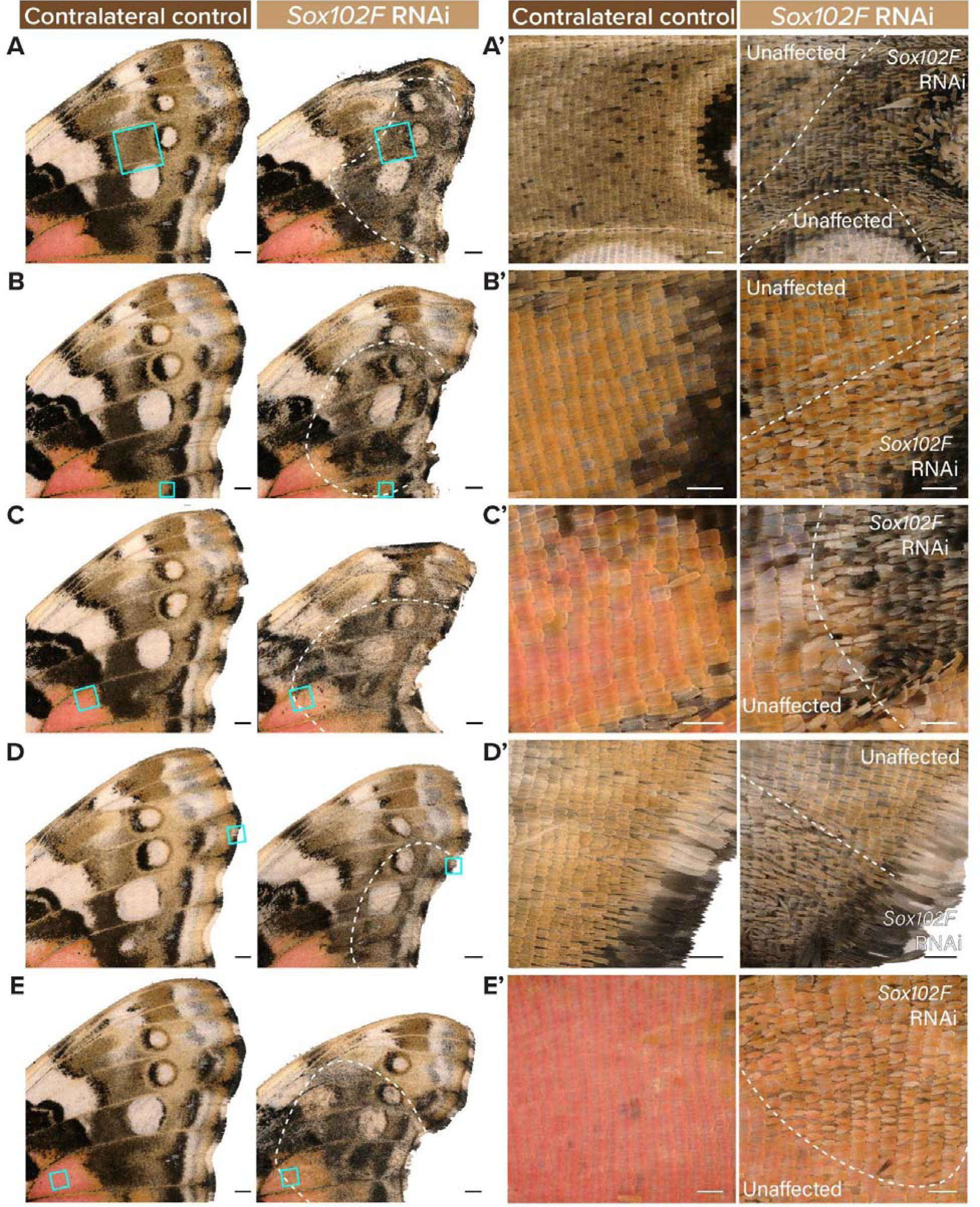
**Representative *Sox102F* RNAi effects on wing scale development, specifically narrower scales independent of colour patterns**. A’-E’ are magnified insets of A-E respectively (cyan boxes indicating origin). Dotted lines demarcate observed RNAi effects. Individual 1 was featured in Figure 4. Scale bars = 1 mm (A-E) and 200 μm (A’-E’).

**Figure S6.**
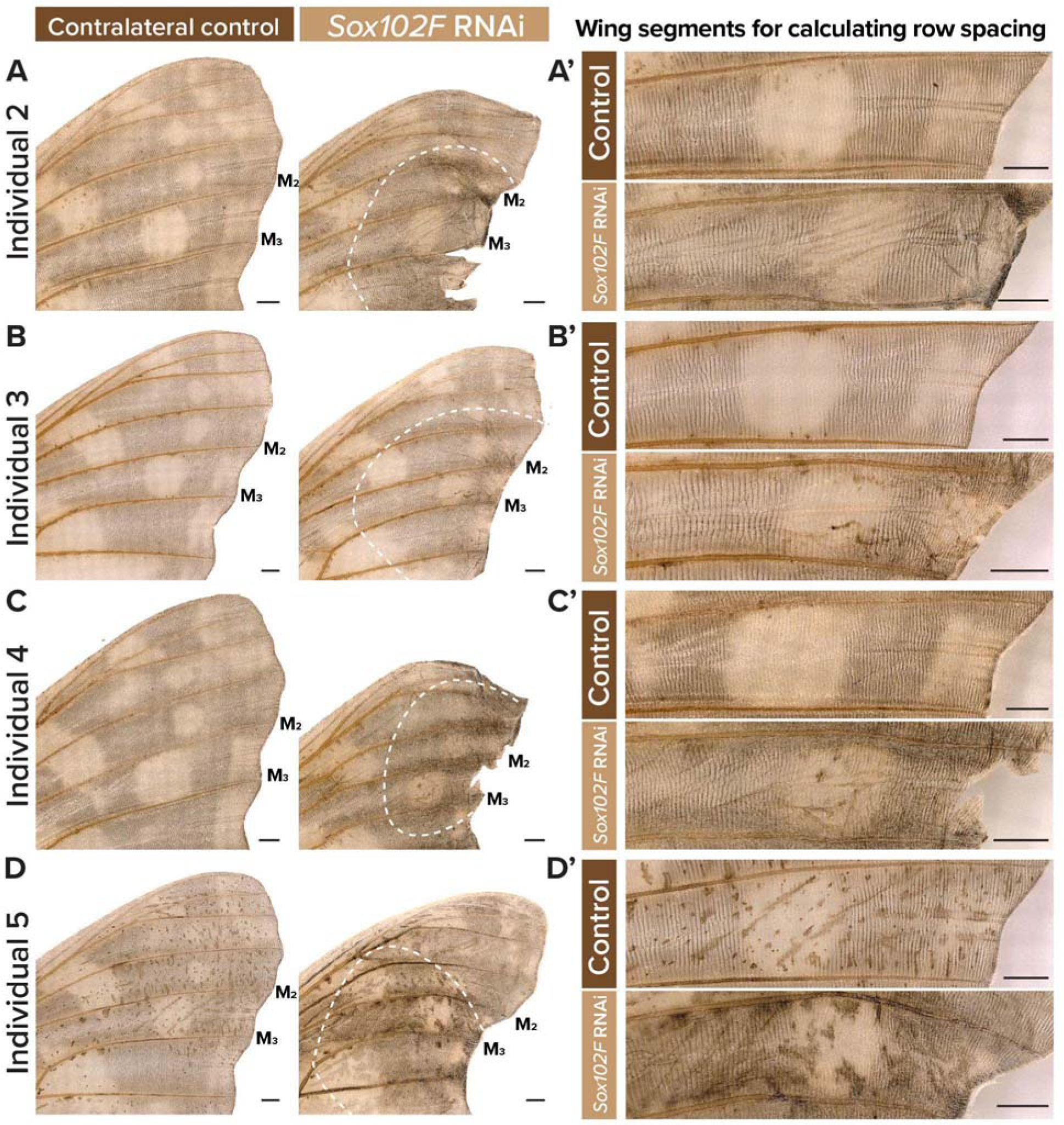
*Sox102F* RNAi treated wings with scales removed. Individuals corresponding to the previous figure. Measurements of row spacing were taken from wing segments flanked by veins M_2_ and M_3_. A’-D’. Dotted lines demarcate observed RNAi effects. Scale bars = 1 mm.

**Figure S7.**
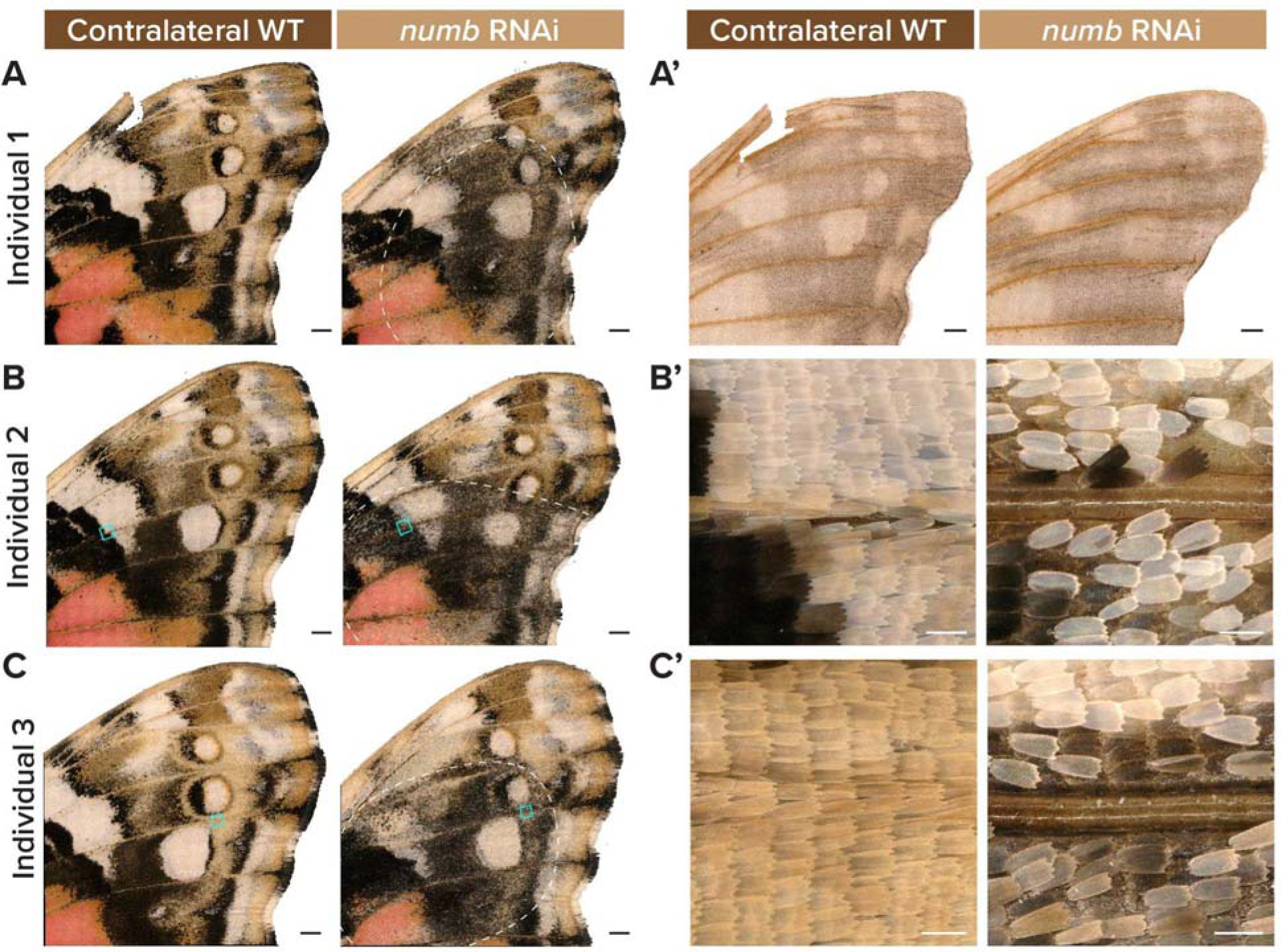
Representative *numb* RNAi effects on wing scale development, specifically the loss of cover scales independent of colour patterns. B’ and C’ are magnified insets of B and C respectively (cyan boxes indicating origin). Dotted lines demarcate observed RNAi effects. Individual 1 was featured in Figure 4. Scale bars = 1 mm (A-A’,B-D) and 100 μm (B’-D’).

**Figure S8.**
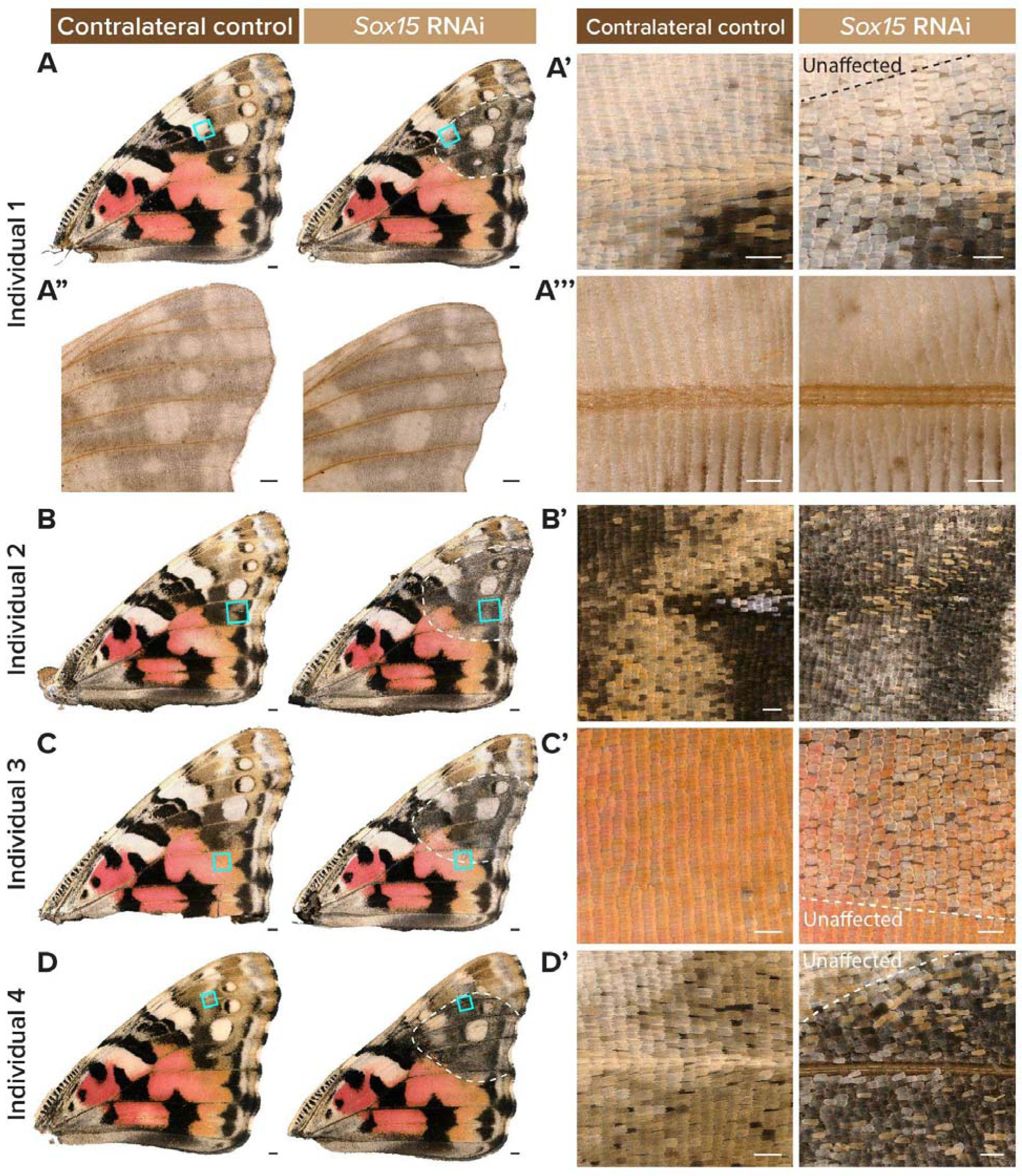
Representative *Sox15* RNAi effects on wing scale development, specifically the loss of cover scales independent of colour patterns. Whole wing (A-D) and magnified views (A-D’) of *sox15* RNAi knockdown wings as compared to their contralateral controls. Dotted lines demarcate observed RNAi effects. Individual 1 was featured in Figure 4. Scale bars = 1 mm (A-D) and 200 μm (A’-D’).

**Figure S9.**
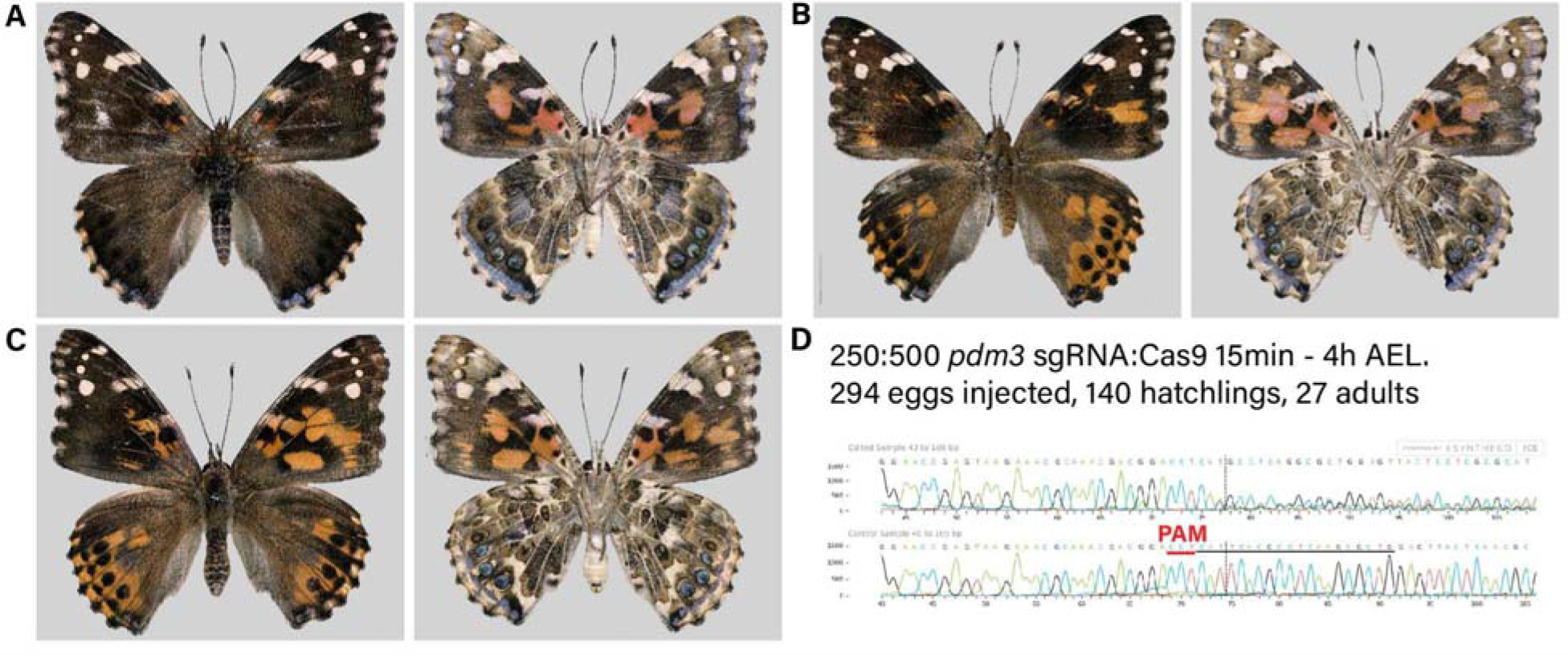
Representative *pdm3* crispants. A-C. *pdm3* mKO results in ectopic black on both surfaces and disruption of pattern elements on ventral surfaces (right). **D.** ICE analysis of crispant for *pdm3* exhibited deletion at sgRNA cut site. Mutant sequence (top) and control sequence (bottom). A was featured in Figure 5.

**Table S1.** Top differentially expressed genes within each cluster in the merged Seurat object.

**Table S2.** Top differentially expressed genes within each cluster in subsetted nuclei within the Seurat object.

**Table S3.** Top 50 genes driving latent time progression inferred from RNA velocity for each of the 3 clusters *SOP*, *scale-* and *socket-building cell*.

**Table S4.**
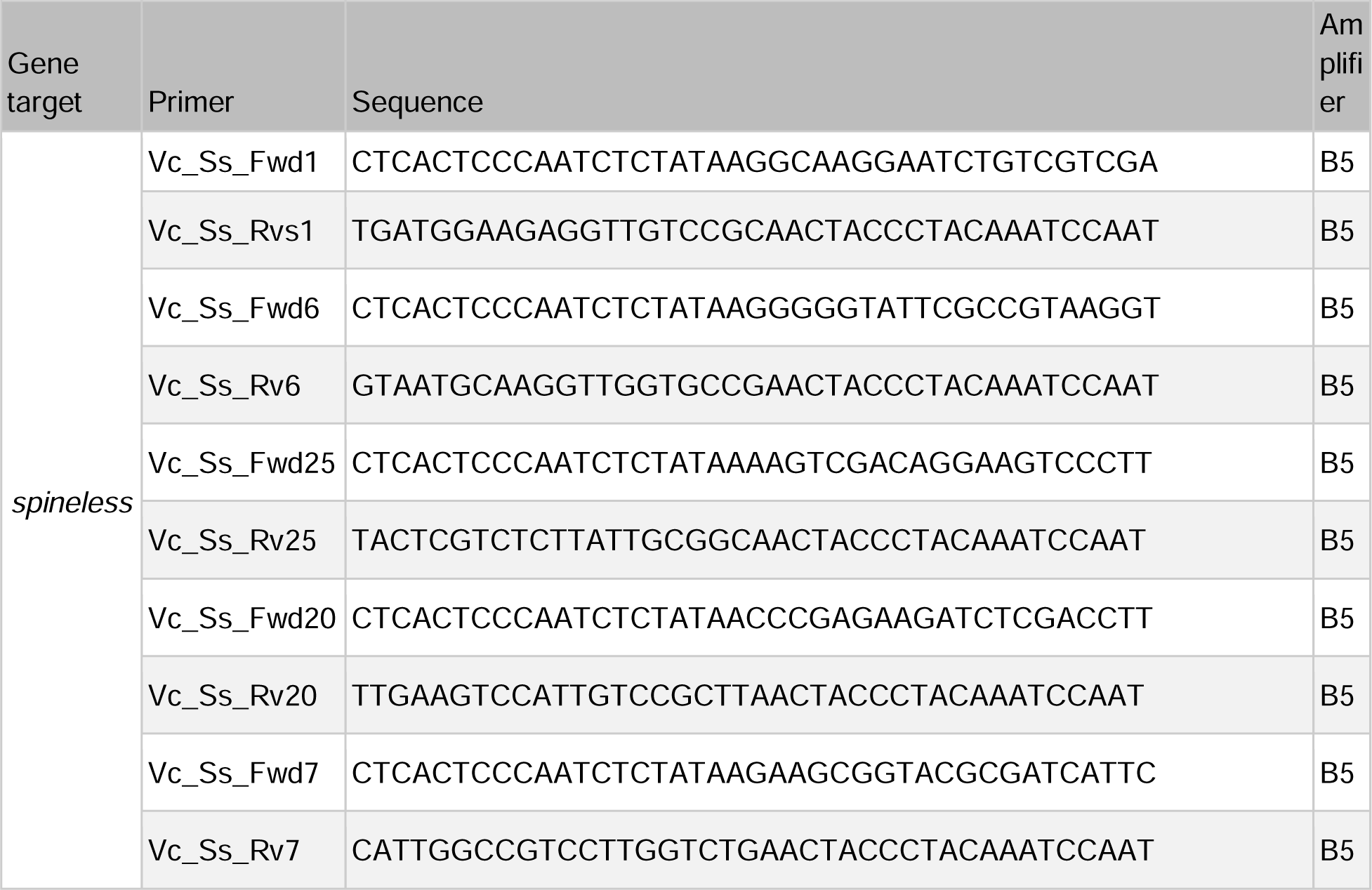
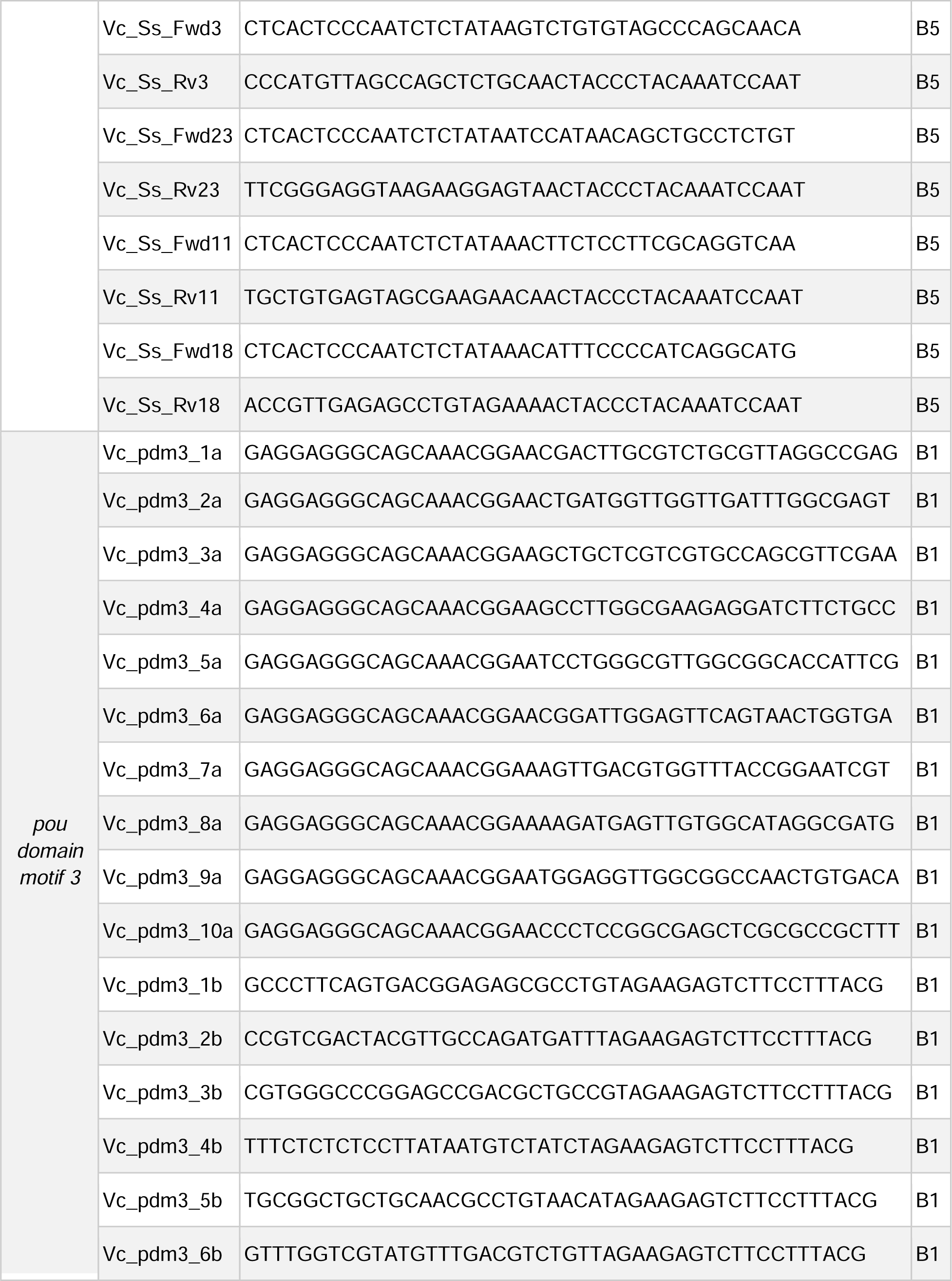
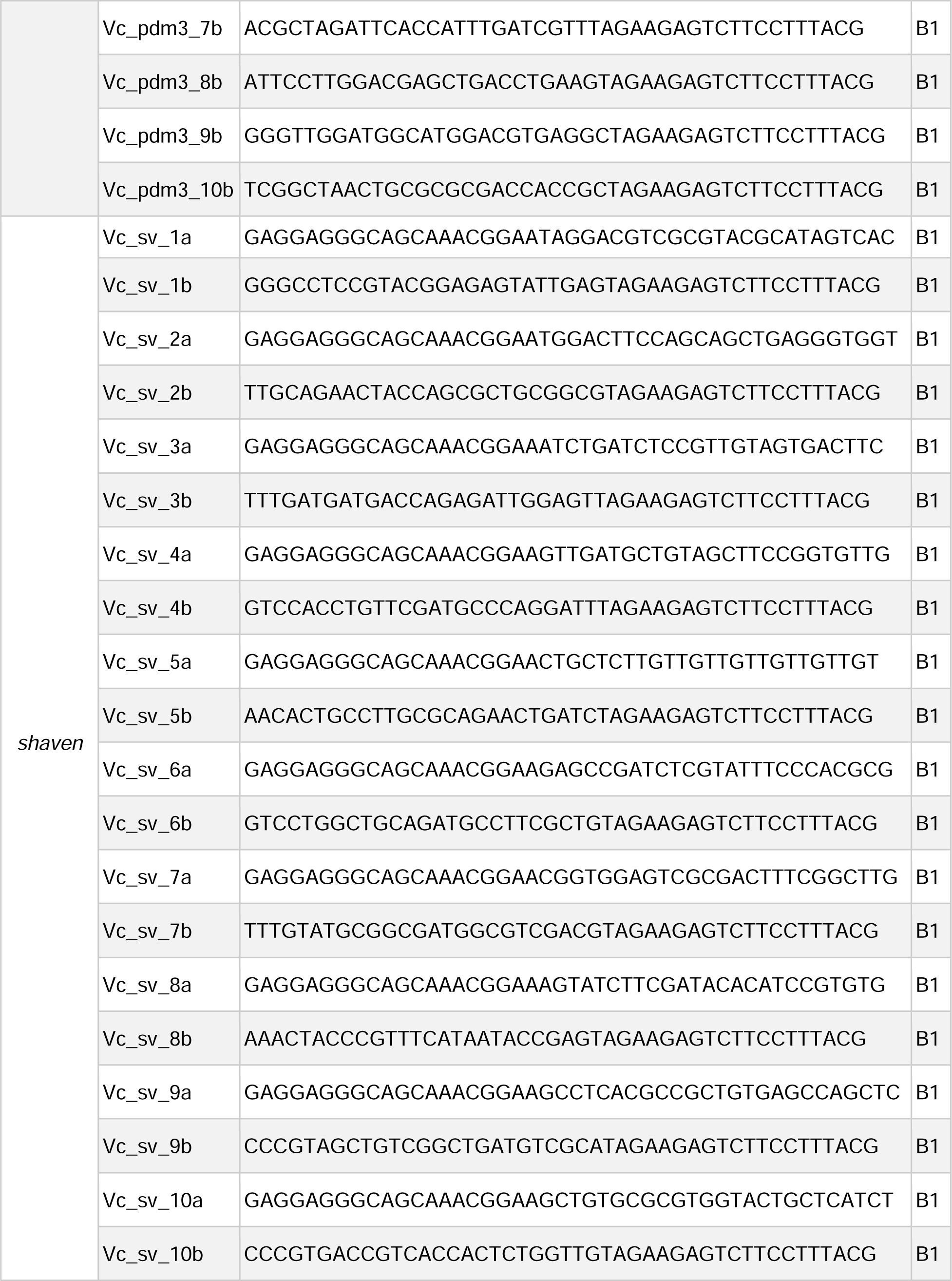
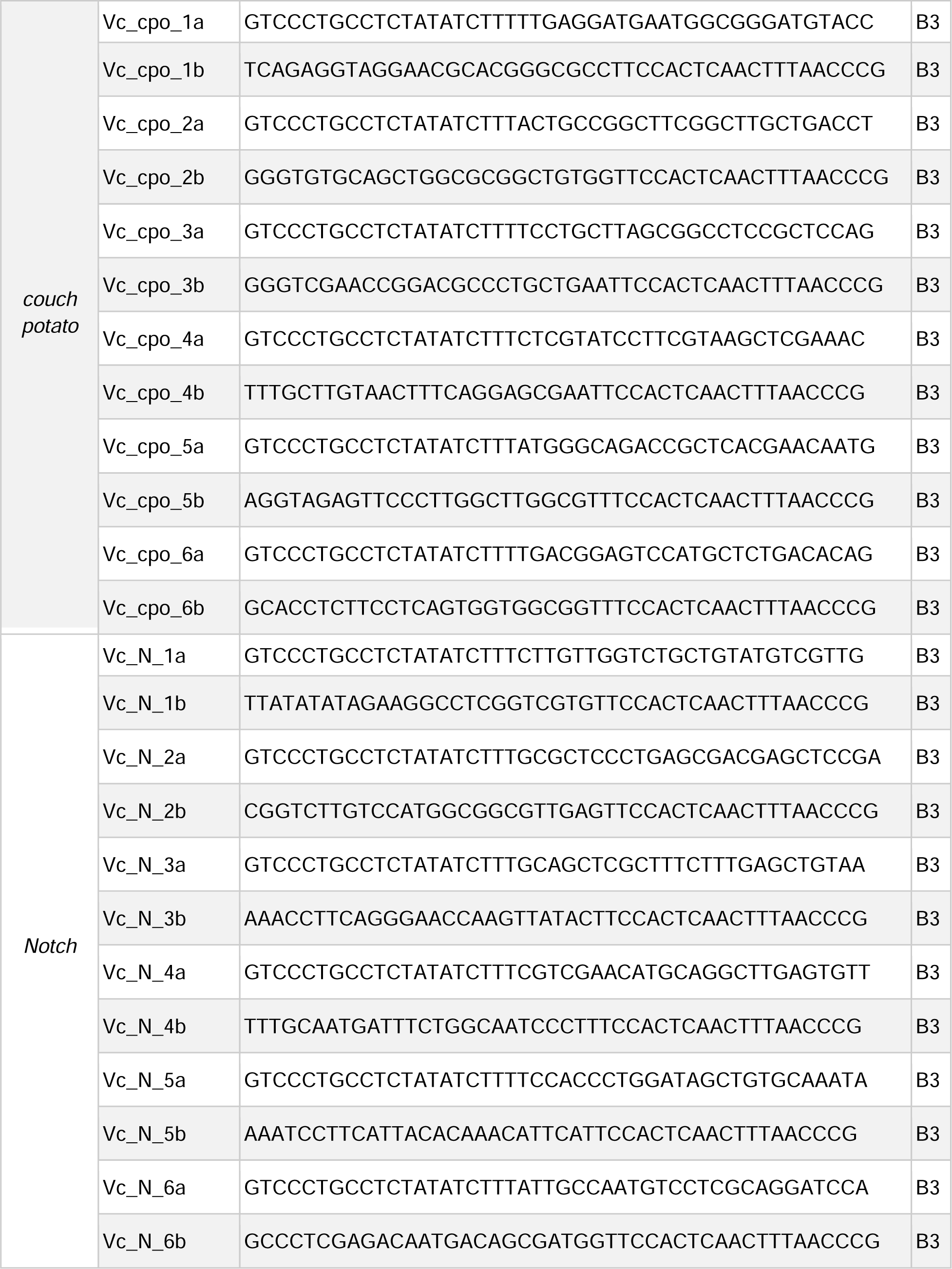
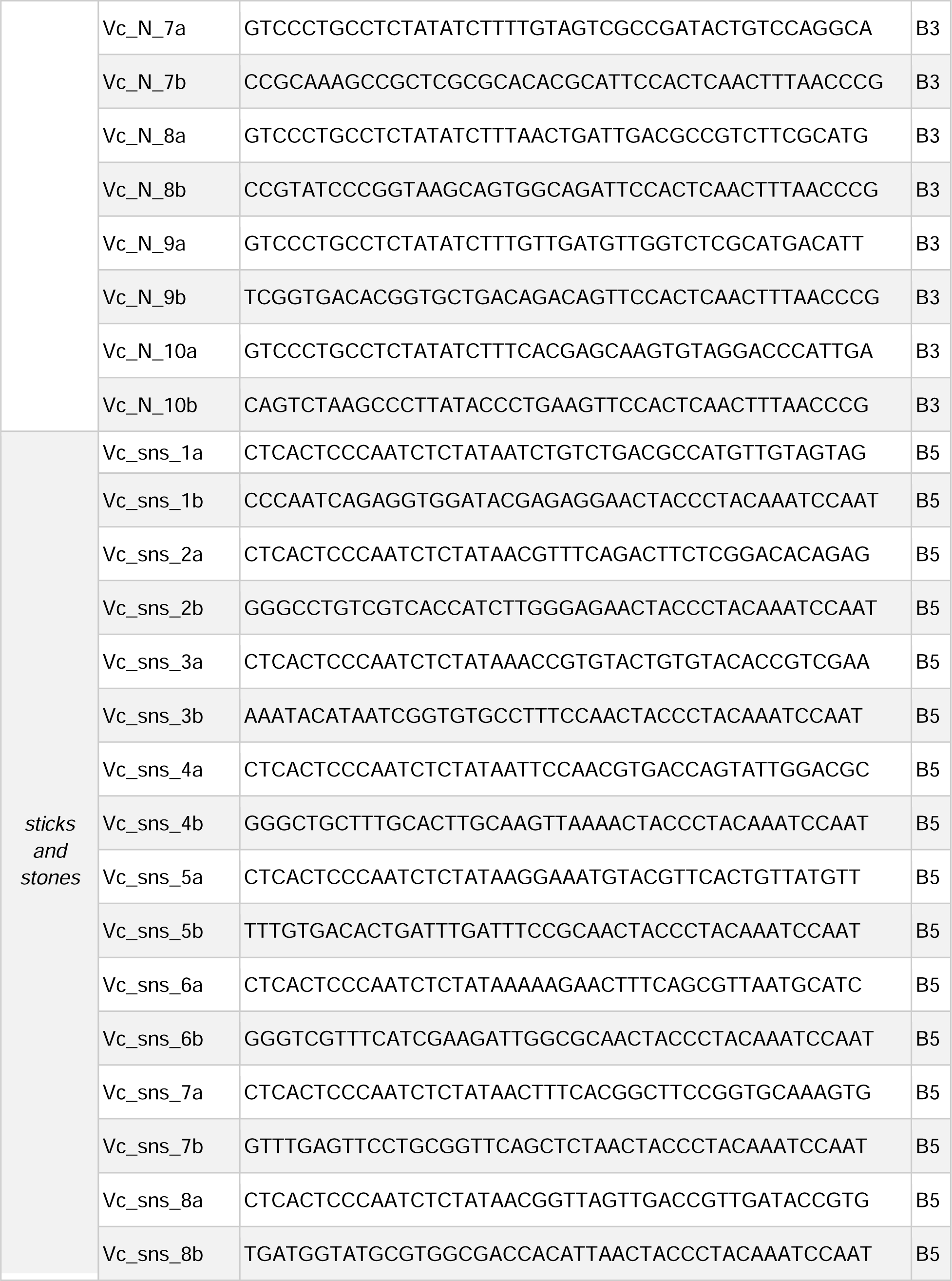
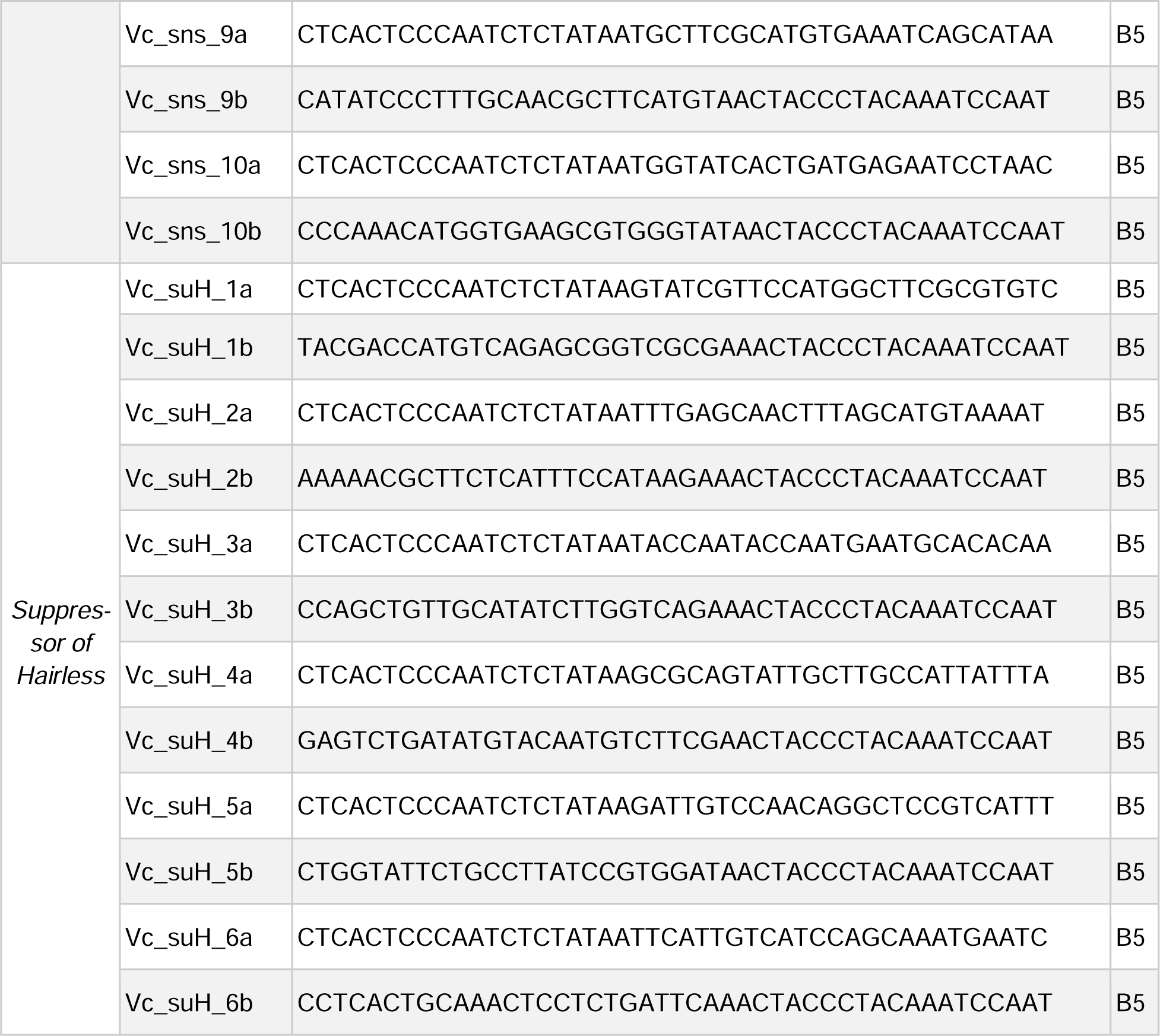
HCR probe sequences used.

**Table S5.**
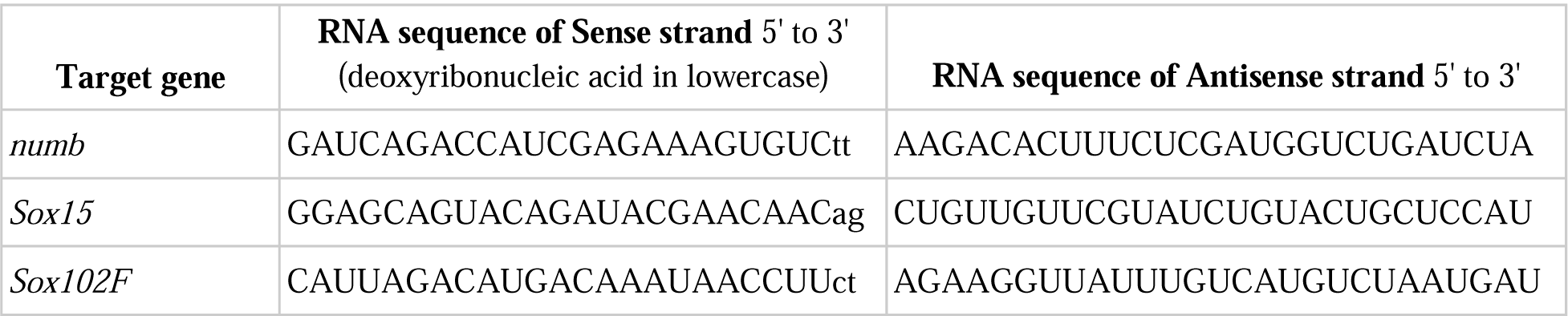
dsiRNA sequences used.

